# FGF signalling plays similar roles in development and regeneration of the skeleton in the brittle star *Amphiura filiformis*

**DOI:** 10.1101/632968

**Authors:** Anna Czarkwiani, David V. Dylus, Luisana Carballo, Paola Oliveri

**Author notes:** Author’s current address: Centre for Regenerative Therapies Dresden, Dresden, Germany (Anna Czarkwiani). Author’s current address: Department Computational Biology & Centre for Integrative Genomics, University of Lausanne, Lausanne, Switzerland (David V. Dylus). Author’s current address: Department of Behavioural Ecology & Evolutionary Genetics, Max Planck Institute for Ornithology, Seewiesen, Germany (Luisana Carballo). These authors contributed equally to this work (Anna Czarkwiani, David V. Dylus). **Corresponding author:** Paola Oliveri.

## Abstract

Regeneration is an adult developmental process considered to be an epiphenomenon of embryonic development. Although several studies have shown that various embryonic genes are expressed during regeneration, there have been no large-scale, direct and functional comparative studies between the development and regeneration of a specific structure in one animal. Here, we use the brittle star *Amphiura filiformis* to characterise the role of the FGF signalling pathway during skeletal development and regeneration. In both processes, we find the ligands expressed in ectodermal cells flanking underlying mesodermal cells, and the receptors expressed specifically by these skeletogenic cells. Perturbation of FGF but not VEGF signalling during skeletogenesis completely inhibited skeleton formation in both embryogenesis and regeneration, without affecting other key developmental processes like cell migration or proliferation. Transcriptome-wide differential analysis identified a highly similar cohort of skeletogenic differentiation genes downstream of the FGF signalling pathway, whereas upstream transcription factors involved in the initial specification of the skeletogenic lineage where unaffected. Comparison to the sea urchin indicated that many of the affected genes are associated with differentiation. Moreover, several genes showed no homology to a cohort from other species, leading to the discovery of brittle star specific, downstream skeletogenic genes. In conclusion, our results show that the FGF pathway is crucial for skeletogenesis in the brittle star, as it is in other deuterostomes, and for the first time provide evidence for the re-deployment of a gene regulatory module during both regeneration and development.

## Introduction

A tempting theory for the evolutionary origins of tissue regeneration suggests it was selected for as a secondary by-product of development and thus shares many similarities with embryogenesis (1,2). In fact, following the unique processes of regeneration (such as wound healing and dedifferentiation), cell specification and differentiation must occur just as they do during embryonic development. In the newt, for instance, gene expression during development and regeneration is conserved. More specifically, the sonic hedgehog gene recapitulates its role in developing limb buds during adult limb regeneration (3); and during elbow joint regeneration in developing chick embryos (4). Developmental genes were often shown to be involved in adult regeneration, although not through a direct comparison of the same animal during embryonic development and regeneration. For example, in planarians many of the components of the genetic network underlying eye development in other species (e.g. *pax6*, *otx*, *six*, *opsin*) have been shown to be expressed and functionally required during adult eye regeneration (5). In salamanders, *Meis* genes under control of the retinoic acid signalling pathway have been shown to be involved in limb regeneration similarly to their role during embryonic limb development (6). Unravelling the function of signalling pathways and transcription factors in development and regeneration can thus shed light on whether adult organisms with the capability of regeneration re-use developmental gene regulatory networks. However, few studies have been carried out thus far, and these mostly compare the expression of a single gene between development and regeneration in the same organism. A new transcriptomic database has recently been published which will provide a platform for whole-transcriptome comparison of development and regeneration in the sea anemone, however no direct functional comparison of specific molecular networks is yet available (7).

Comparing the role of signalling pathways in embryogenesis and regeneration provides a compelling strategy to understand the extent of similarities between gene regulatory networks driving these two developmental processes. A good example of this is the fibroblast growth factor (FGF) signalling pathway, which has been implicated in a wide range of biological processes during development such as cell migration, differentiation and proliferation, during both wound healing and regeneration (8,9). FGF signalling is highly conserved among metazoans, although varies substantially with respect to the number of its ligands in different species. Mesoderm formation is one of the highly conserved processes for which the role of FGF signalling is employed similarly across the Bilateria, including vertebrates (10,11), hemichordates (12) sea urchins (13), flies (14) and beetles (15). Additionally, this pathway is also heavily involved in post-embryonic developmental processes. Regeneration in zebrafish, *Xenopus* and salamanders relies on the expression of FGF genes in the blastema, and applying FGFR inhibitors results in regenerative defects (16–20). The FGF signalling pathway also plays important roles in development and regeneration of the vertebrate skeleton. Mutations in both ligands and receptors were found to cause a variety of congenital disorders including craniosyntoses, chondrodysplasia (21–23), and multiple types of gross skeletal development abnormalities in mouse models and humans (24). Similarly, multiple FGFs and FGFRs are expressed during fracture healing and bone regeneration (25). Importantly, the precise roles and effects of FGF inhibition during postembryonic morphogenesis is not well understood (26).

The role of FGF signalling in skeletogenesis also extends to the invertebrate deuterostomes, namely echinoderms, which are an excellent experimental system for studying the gene regulatory networks (GRN) underlying development (27–30). It has been shown that FGF signalling is necessary for guiding skeletogenic mesenchymal cell migration and formation of the embryonic skeleton in the sea urchin *Paracentrotus lividus* (31). Interestingly, in a different species, *Lytechinus variegatus*, FGF inhibition using an *fgfa* morpholino (also called *fgf9/16/20*) produces a much milder phenotype, whereby the mesenchymal cells migrate normally and the embryos form shortened skeletal rods (32). In addition to FGF signalling, the VEGF signalling pathway is also involved in skeletogenesis in both species. Perturbation of the *vegf3* ligand in the sea urchin interferes with both correct skeletogenic cell migration and skeletal rod formation (32–34). It seems clear that both of these pathways have essential, often interconnected and non-redundant roles in skeletogenesis in the sea urchin embryo. It is not well understood, however, whether these pathways regulate different downstream effector genes, and whether their role is conserved during adult skeletogenesis in echinoderms.

Recently, several studies have established the brittle star *Amphiura filiformis* (*Afi*) as an experimental system for skeleton formation in both embryonic development (35,36) and adult regeneration (37,38). Characterization of these processes showed that the skeletogenic cells of both the adult and embryo are mesenchymal and express an array of skeletogenic specification transcription factors such as *alx1*, a gene belonging to a family of transcription factors with a conserved role in skeleton development in echinoderms (39), and vertebrates (40,41). Adult skeletogenic cells also express downstream embryonic skeletal differentiation genes, including *c-lectin, p58b, p19* and *α-coll* (35,37,38). Moreover, transcriptomic data for both the embryonic stages (36,42) and the adult regenerating and non-regenerating arms (43,44) are now available. With this wealth of information on regeneration and early development of the skeleton, this species can be used to directly compare the role of FGF signalling in both processes within the same animal at different life stages.

In this study, we carry out for the first time a large-scale, side-by-side comparison of the development and regeneration of a skeletogenic structure in embryos and adults of the same species in the context of FGF signalling. We first characterize the expression of FGF signalling components during embryogenesis and adult arm regeneration. We then use an FGF signalling inhibitor (SU5402) in embryos and adult *A. filiformis* to determine the effect of disrupting this pathway. We find that perturbation of FGF signalling in brittle stars results in failure to form skeletal spicules in both the embryos and in the regenerating arms. Using an unbiased systems approach comparing control and treated embryos, we find several brittle star specific skeletogenic genes. Moreover, many of these are affected similarly in embryos and in adult regenerating arms, suggesting a conservation of pathway components and network connections between these two processes. Additionally, we did not find any upstream genes affected by the perturbation of the FGF signalling pathway in the skeletogenic cells. Ultimately, our study provides the first direct evidence for an analogous role of FGF signalling in skeletogenesis between embryonic development and adult regeneration in the same species working downstream from the specification tier of the skeletogenic GRN.

## Results

### Evolutionary relationships of FGF and VEGF signalling components in echinoderms

To characterize the *fgf* and *vegf* signalling genes, we first surveyed an embryonic transcriptome encompassing the entirety of development (from cleavage stage to pluteus larvae) (36), and transcriptomes from adult regenerating and non-regenerating arms (43) of the brittle star *A. filiformis* for potential homologs. To do this, we combined a BLAST search using selected candidates (e-value 1e-6) from the sea urchin database (45) with a hidden Markov model search against PFAM domains of both ligands and receptors (46). Using this strategy, we found 2 Fgf ligands and 3 Fgf receptors, and 2 Vegf ligands and 1 Vegf receptor in *A. filiformis* (S1 Table).

To better understand the evolutionary relationships of our *A. filiformis* genes relative to chordate signalling systems, we computed five phylogenetic trees using the amino acid sequences of candidates from 41 species spanning all major clades of echinoderms, chordates (e.g. mouse, rat etc.) and non-deuterostome outgroups species such as the pacific oyster (*Crassostrea gigas*). We observe that two Fgf ligands are placed in an echinoderm group sister to their respectively independently duplicated genes in chordates (S1A and S1B Fig), Afi-Fgf9/16/20 and Afi-Fgf8/17/18, suggesting a duplication of the FGF ligands at the base of the deuterostome lineage. Different evolutionary relationships are revealed for the Fgf receptors. The Afi-Fgfr1 to Afi-Fgfr2 and Afi-Tk9 are all in a group with other echinoderm FGF receptors and sister to the chordate FGFR1, FGFR2 and FGFR4 receptors. Hence, suggesting that gene independent duplication events from a common ancestral FGF receptor occurred in chordates as well as in echinoderms (S1C Fig). Concerning the Vegf ligands, we also observe independent duplication events in chordates (VEGFA and VEGFB) as well as echinoderms (Vegf3 and Vegf4) (S1D Fig). The only VEGF receptor of echinoderms forms a sister group to three VEGF receptor genes in chordates (S1E Fig). Due to our broad sampling of echinoderm species we were able to place genes from other echinoderms and hemichordates in the same grouping, which confirm and strengthen the evolutionary relations of the *A. filiformis* genes. Here, however, we specifically focus on the genes that show clear orthology between the sea urchin and brittle star, as the sea urchin has a well-annotated genome. Ultimately, this analysis allows us to bring our results into a broader evolutionary context when comparing across different species.

### FGF signalling genes are expressed during both embryonic development and adult arm regeneration

To better understand the role of *fgf* signalling genes in the context of skeletogenesis, we first analysed the expression of ligands and receptors during embryogenesis and adult regeneration using *in situ* hybridization (ISH) and NanoString for transcript quantification. For this purpose, we selected the most likely corresponding stages between development and regeneration based on morphological features and gene activity (S2 Fig). Specifically, we focused on the developmental stage when the skeletogenic lineage is segregated from other mesodermal cells and specific skeletogenic genes are expressed (blastula and mesenchyme blastula stages of development Fig 1A; stages 3-5 during arm regeneration Fig 1B) and when skeletal spicules appear (gastrula stage of development; Fig 1A; stage 3-5 during adult arm regeneration; Fig 1B) (35–38).

**Figure 1:**
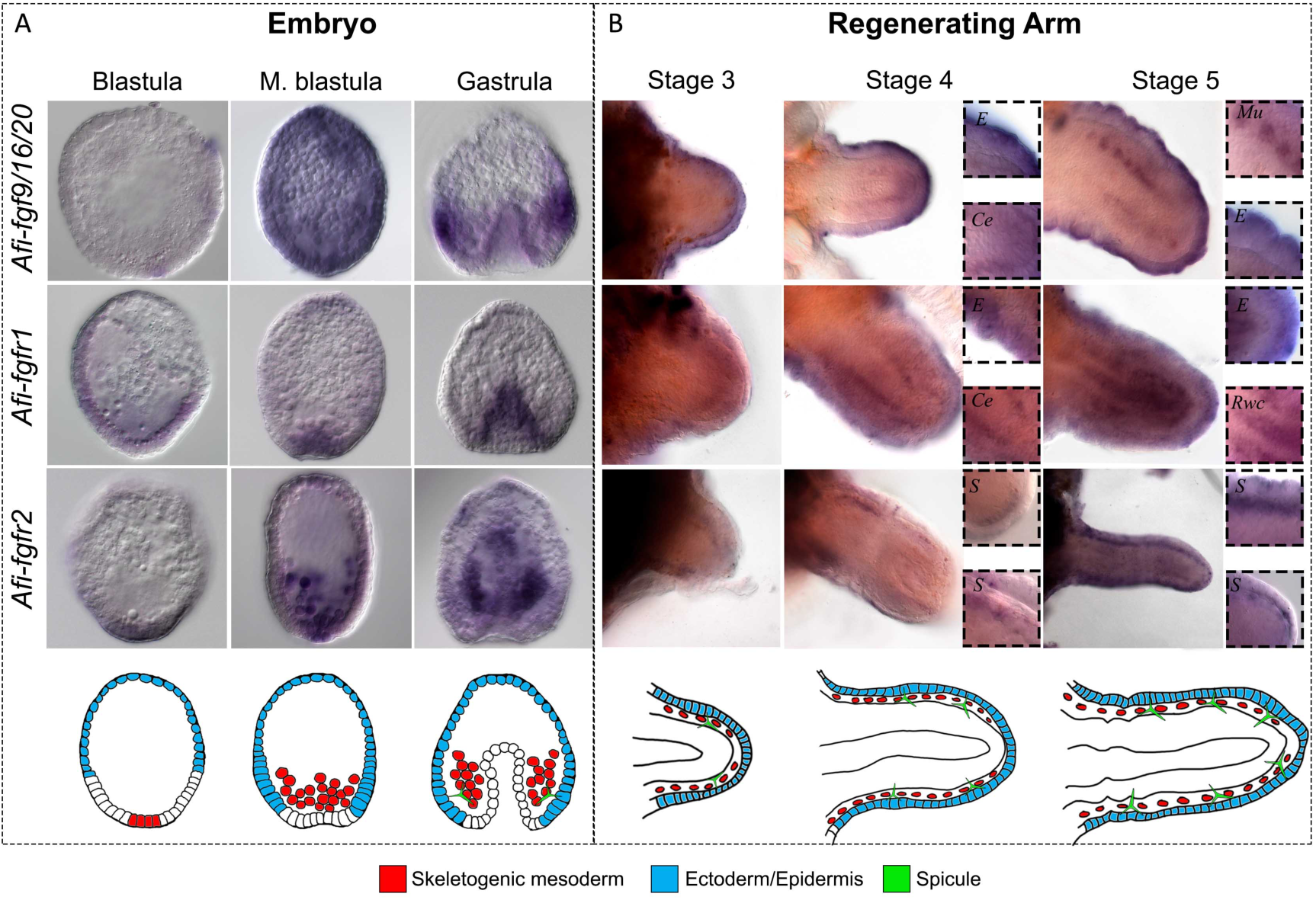
Expression of FGF signalling components in embryos and early regenerating arm stages of *A. filiformis*. A) Top: WMISH on embryos at blastula, mesenchyme blastula and gastrula stages of development showing the expression of *Afi-fgf9/16/20, Afi-fgfr1, Afi-fgfr2*. Bottom: schematic diagram of major relevant cellular domains in blastula, mesenchyme blastula and gastrula stage embryos of A. filiformis. B) Top: WMISH on regenerates at stages 3, 4 and 5 showing the expression of *Afi-fgf9/16/20, Afi-fgfr1, Afi-fgfr2*. Insets show detail of expression patterns. Bottom: Schematic diagram of major relevant cellular domains at stages 3, 4 and 5 of early arm regeneration in *A. filiformis*. E –epidermis, Ce –coelomic epithelium, Mu –metameric units, Rwc -radial water canal, S –skeletogenic cells in dermal layer. Scale bars: 100µm.

We characterized the expression of the FGF signalling pathway components of these two developmental processes. In the embryo, the *Afi-fgf9/16/20* ligand is first detectable at mesenchyme blastula stage, between 15 and 18 hours post fertilization (hpf) ubiquitously (Fig 1A; S2A Fig). It then becomes confined to a band in the ectodermal domain at the boundary of the endoderm, with higher expression in two domains adjacent to the clusters of mesenchymal cells that will produce the skeleton of the embryo at gastrula stage (Fig 1A; S2A Fig). During regeneration, *Afi*-*fgf9/16/20* is expressed in the epidermis throughout early stages (stage 3-5; Fig 1B; S2D Fig) and expands to an additional domain adjacent to the radial water canal in patches of cells that most likely correspond to the newly forming metameric units of the regenerating arm at stage 5 (Fig 1B; S2D Fig). During development, the receptor *Afi-fgfr1* is expressed in the vegetal half of the embryo at blastula stage, and endoderm and non-skeletogenic mesoderm from mesenchyme blastula stage to gastrula stage (Fig 1A; S2B Fig). It also exhibits a highly dynamic expression during adult arm regeneration in several territories including the epidermis, coelomic epithelium and radial water canal (Fig 2B; S2E Fig). Conversely, the receptor *Afi-fgfr2* is first specifically expressed in the skeletogenic mesoderm (SM) cells at mesenchyme blastula stage and expands to the non-skeletogenic mesoderm at gastrula stage during embryonic development (Fig 1A; S2C Fig). During stages 4 and 5 of regeneration it is expressed in the dermal layer where the skeleton first appears during regeneration (29; Fig 1B; S2F Fig). Expression of FGF signalling pathway components at late stages of regeneration persist in similar territories (epidermis for the *fgf9/16/20* ligand and skeletal domains for *fgfr2* receptor; S3 Fig). Interestingly, the expression of *fgf9/16/20* in the ectoderm and the *fgfr2* receptor in mesenchymal cells is comparable to the expression of their orthologs in sea urchin development (31). With respect to the organization of mesenchymal cells and their proximity to the ligand-expressing ectodermal domain cells, sea urchin and brittle star embryos share a similar topology. The highly comparable expression of FGF signalling pathway components between brittle stars and sea urchins (31,47), as well as during development and regeneration suggests a conservation of the mesoderm-ectoderm interaction necessary for skeletal formation.

**Figure 2:**
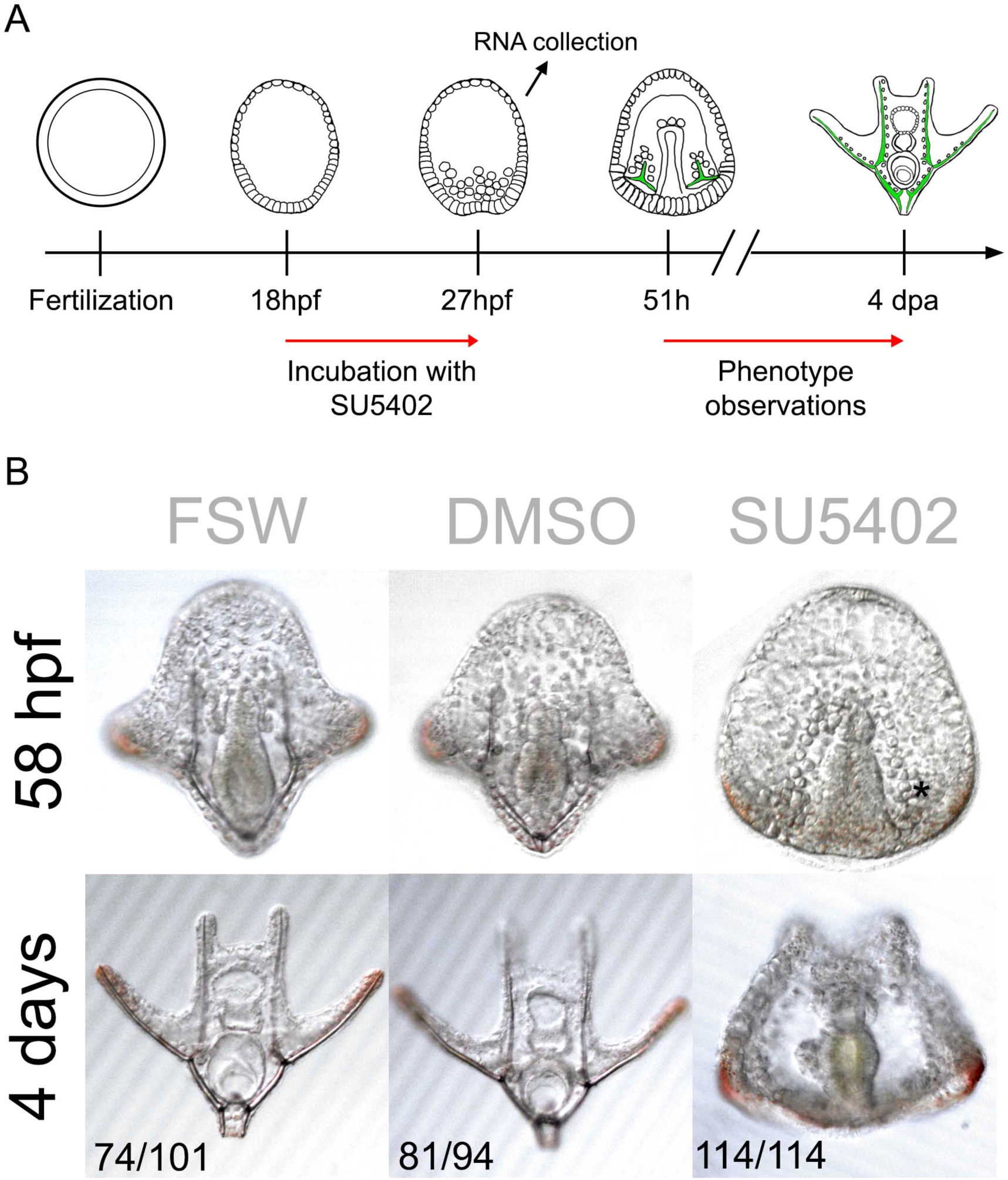
FGF signalling perturbation using the SU5402 inhibitor in brittle star embryos. A) Schematic diagram of experimental procedure for SU5402 treatment. B) Phenotypic analysis of SU5402-treated *A. filiformis* embryos and controls at 58hpf and 4 days post fertilization shows that perturbation of FGF signalling results in embryos with no skeletal spicules forming. Numbers at the bottom show counts for embryos observed with the represented phenotype/total embryos counted.

### FGF signalling perturbation with SU5402 inhibits skeleton formation in both embryos and adult regenerating arms

To analyse the role of FGF signalling in skeletogenesis during brittle star embryonic development and adult arm regeneration we applied the SU5402 inhibitor, a small molecule well-known to specifically inhibit the function of FGFRs by competing with ATP for the binding site of the catalytic domain of tyrosine kinase (48). This inhibitor has been successfully used to disrupt FGF signalling during both embryogenesis and regeneration (18,49–51) in many organisms.

For developmental characterization, we treated brittle star embryos with 10µM SU5402, alongside non-treated filtered seawater (FSW) and DMSO (used as the solvent for the drug) controls at 18hpf (hours post-fertilization) preceding skeletogenic mesoderm ingression (35). This time point was used to avoid interfering with potential early functions of FGF signalling during cleavage stages and to specifically focus on skeletogenesis (as this also corresponds to the temporal onset of *fgfr2* expression in skeletogenic cells, see Fig 1A and S2C Fig). At 27hpf we collected the embryos for RNAseq and NanoString analysis (Fig 2A) to assess the early response to signalling inhibition and to avoid secondary effects of FGF perturbation for a prolonged period. At this stage the treated embryos are indistinguishable from controls showing a timely ingression of the primary mesenchymal cells. Subsequently, we scored several embryos at late gastrula and pluteus stages for the formation of spicules. All SU5402-treated embryos failed to develop skeletal spicules (100%, n=114), compared to 0.2% DMSO (13.8%, n=94) and FSW controls (26.7%, n=101) (Fig 2B). Despite having no visible defects in skeletogenic mesoderm ingression, archenteron invagination or overall survival (58 hpf; Fig 2B), the perturbed embryos did not develop a skeleton, even at late stages of development (4 dpf; Fig 2B).

A similar treatment was performed in regenerating arms to functionally assess the role of FGF signalling during adult skeleton regeneration. We applied the SU5402 inhibitor to amputated arm explants, which can survive separated from the main body for several weeks and continue to regenerate properly (52). The explants were incubated in 10µM SU5402 from stage 2 (prior to formation of skeletal spicules; (38)) for 24h after which they were scored for phenotype and collected for further analyses (Fig 3A). The development of biomineralized skeletal primordia (or spicules) can be monitored by incorporation of calcein, a green fluorescent dye that labels the newly deposited CaCO_3_ (53). FGF signalling perturbation using this method caused inhibition of skeletal spicule formation in the majority of arms (78.1%, n=41), as shown by the absence of calcein staining in the dermal layer, compared with 0.1% DMSO controls (7.7%, n=39) and non-treated FSW controls (8.1%, n=37) (Fig 3B). All arm explants were alive and mobile after treatment (S1 Movie), however only the DMSO and FSW controls continued to regenerate 48h after treatment (S4A Fig). Since treated explants did not elongate, and to rule out possible toxic side effects, we examined whether the explants retained cell proliferation ability. Interestingly, even though SU5402 treated explants failed to regenerate further (n=8; S4A Fig) we found that cell proliferation was not affected by the inhibition of FGF signalling (n=4; S4B and C Fig). This provided evidence that not all cellular mechanisms have been affected by the treatment, but rather a specific effect has been exerted on regeneration of different tissues, including the skeleton. Importantly, the application of SU5402 led to a reduction of skeleton during both embryonic development and regeneration.

**Figure 3:**
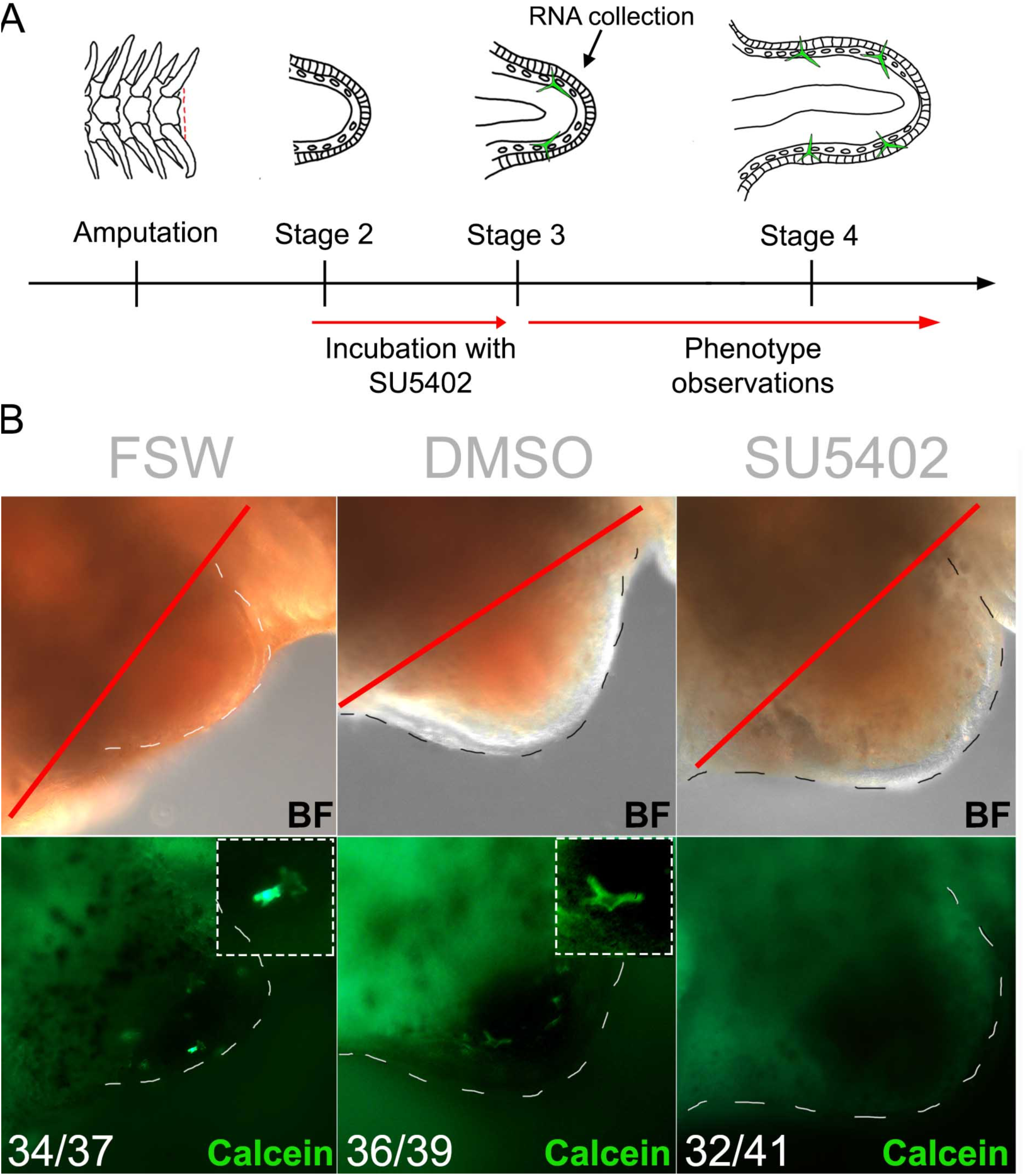
FGF signalling perturbation using the SU5402 inhibitor in brittle star regenerating arm explants. A) Schematic diagram of experimental procedure for SU5402 treatment. B) Phenotypic analysis of SU5402-treated *A. filiformis* regenerates and controls at 24 hours post treatment (stage 3) shows that perturbation of FGF signalling inhibits spicule formation. Numbers at the bottom show counts for explants observed with the represented phenotype/total explants counted. Red line –amputation plane. Dashed lines –outline of regenerating bud.

### VEGF signalling perturbation with Axitinib mildly inhibits skeleton formation in both embryos and adult regenerating arms of A. filiformis

The VEGF signalling pathway plays a crucial role in sea urchin skeletogenesis and the expression of its ligand in the ectoderm and of its receptor in the mesenchymal cells resembles the expression of components of the FGF pathway (54). SU5402 has been shown to have some mild inhibitory effects on this pathway at high concentrations: 50μM and above (55), therefore to determine to what extent inhibition of the VEGF pathway alone could interfere with skeletogenesis, we characterized the expression of its components (S5A and B Fig) and inhibited it with a VEGF specific inhibitor: Axitinib, which selectively inhibits VEGF receptors by blocking their cellular autophosphorylation (56) (S5C and D Fig). Interestingly, although the expression patterns of VEGF ligands and receptors is strikingly similar to FGF components in embryos and regenerates in the brittle star (S5A and B Fig), inhibition of the VEGF signalling pathway using the pharmacological agent Axitinib resulted in a much milder phenotype in respect to skeleton development in the embryos and regenerating explants of *A. filiformis* compared with the phenotype obtained with the SU5402 treatment (compare Fig 2 and 3 with S5C and D Fig). Axitinib treated embryos usually formed one spicule during early development and this spicule elongated but failed to be patterned correctly (n=89/118) compared with normal skeletogenesis in FSW (n=102/119) and DMSO controls (n=101/123) (S5C Fig). Only 36.6% of treated explants (n=41) showed reduced or absent spicules compared to 13.6% in DMSO controls (n=44) and 10% in FSW controls (n=40) (S5D Fig). At the concentration used in this study, it is unlikely that SU5402 inhibition impinges significantly on the VEGF pathway, and specific inhibition of VEGF signalling shows that it is not strictly required for biomineralization to occur, but most likely for further patterning of the skeletal elements. We thus only focused on the molecular network affected by SU5402 treatment from this point on.

### Genes downstream of the FGF signalling pathway are involved in embryonic skeletogenesis

To identify potential genes involved in skeletogenesis and other unknown targets of the FGF signalling pathway, we conducted a transcriptome-wide analysis of SU5402-treated embryos relative to controls (Fig 4). In this analysis we used a log fold change threshold of ±1.6 log2(SU5402/DMSO), as used for sea urchin (57,58), to select up- or down-regulated candidates in the transcriptome dataset (59). With this threshold, we obtained 140 downregulated and 2,366 upregulated transcripts (Fig 4A). As SU5402 inhibited skeleton development (Fig 2B), we focused our attention on the downregulated genes to pinpoint potential candidates that may be involved in skeleton formation. In the 140 downregulated transcripts, and using our transcriptome annotation (36), we found 101 sea urchin homologs of which only three were TFs (transcription factors; *Afi-six1/2, Afi-egr* and *Afi-soxD1*) and 16 were known skeletogenic genes (Fig 4C). To improve the power of our predictions and to validate the differential transcriptome analysis, we performed NanoString on 123 selected candidates (26/140 downregulated, 5/2366 upregulated and other genes potentially involved in regeneration and development of skeleton; S2 Table) on two biological replicates, and QPCR on three biological replicates using a subset of these 123 candidates. In order to compare data across different technologies, quantitative data were collected on the same sample using all three technologies (RNA-seq, QPCR and NanoString) and used to identify conversion factors to bring all data from different biological replicates on a comparable quantitative scale (details in Methods). With this approach we were also able to compute additional significance values. 24 genes showed significant differences from 0 (p*<0.05) of which 12 were below log2(fc) −1.6 and 3 above +1.6, and the residual 9 were close to ±1.6 (Fig 4C and S11 Fig). Interestingly, in the embryo only a few transcription factors were affected in the transcriptome-wide analysis and none of them are known TF expressed in the SM (skeletogenic mesoderm). On the contrary, FGF and VEGF signalling components showed significant differential expression: both the two receptors specifically expressed in SM cells (*Afi-fgfr2* and *Afi-vegfR*) are down regulated in SU5402 treated embryos, while the *Afi-vegf2* ligand is strongly upregulated. To address the spatial expression of differentially expressed transcripts, we performed WMISH on selected genes from another biological replicate (Fig 4B). WMISH on four downregulated transcripts specifically expressed in SM cells (*Afi-msp130L, Afi-tetraspanin*, *Afi-tr9107, Afi-slc4a10)*, and one ectodermally expressed gene *Afi-egr*, consistently showed loss of expression in SU5402-treated samples, whereas an increase in ubiquitous expression of *Afi-alx/arx* was detected, a gene identified as upregulated in all our quantitative expression assays. As a negative control the unaffected gene *Afi-*α*coll* shows no change of expression (Fig 4B). These data indicate that the combination of different technologies on different biological replicates resulted in a reliable list of candidate genes that were affected by SU5402. Moreover, the observed effect on skeletal downstream genes, such as *msp130*, *slc4a10, kirrell, p-58* and more, rather than transcriptional regulators, suggest a role of FGF signalling primarily in the differentiation step of skeleton development rather than in specification. Interestingly, components of both FGF and VEGF signalling are also affected in SU5402 suggesting a complex crosstalk between these two signalling pathways in the development of *A. filiformis* skeleton.

**Figure 4:**
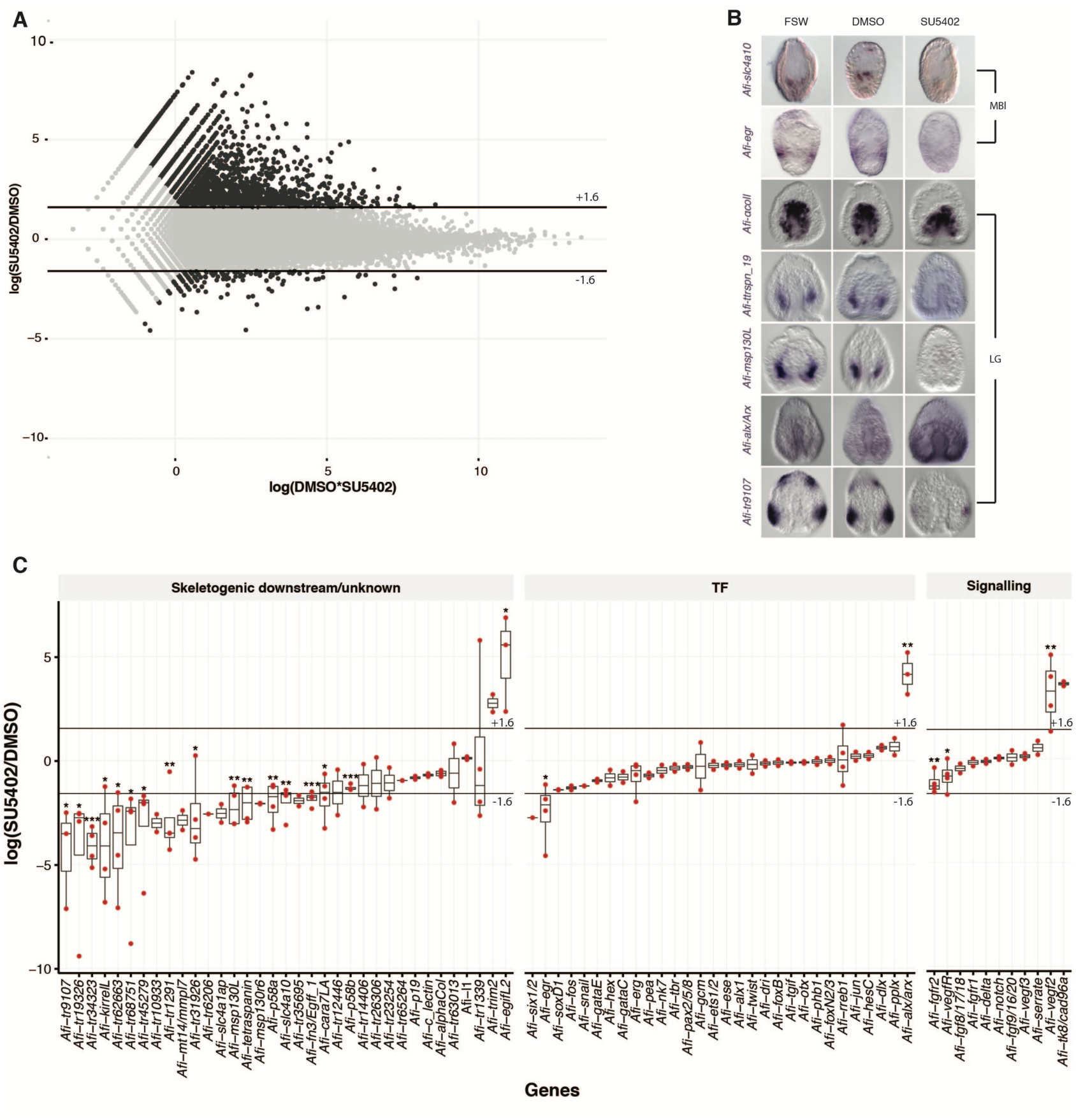
Differential analysis for SU5402 treated samples and WMISH analysis on treated embryos. (A) MA-plot showing upregulated samples on top and downregulated samples on the bottom. (B) WMISH on embryos treated with SU5402 that were fixed at gastrula stage. *Afi-αcoll* was used as negative control and no change in expression is observed. *Afi-ttrspn 19, Afi-msp130L*, and *Afi-tr9107* are downregulated and *Afi-alx/arx* is upregulated in SU5402 treated samples. C) Box plot summarizing differential gene expression in embryos treated with SU5402 and DMSO showing consistency between transcriptome, qPCR and nanostring quantification strategies represented as log2(DMSO/SU5402).

### Genes downstream of the FGF signalling pathway are also involved in skeleton regeneration

To compare the genes transcriptionally regulated by FGF signalling between embryonic development and adult regeneration we performed a large-scale analysis of the effects of SU5402 perturbation in explants using NanoString. We used a code set of 123 genes and quantified 3 biological replicates of RNA extracted from 10 individual arm explants treated with SU5402 for 24h (at stage 2) relative to controls. To detect differentially expressed candidates a log fold change of 1 was used as a threshold of significance, similarly to previously published work (60). In this analysis we found 25 differentially expressed genes (10 upregulated and 15 downregulated, discussed further in next section). We performed WMISH on at least three SU5402-treated explants and relative controls for each gene from a selected group of transcripts (S7 Fig) to validate our quantitative analysis (Fig 5). Transcripts classified as downregulated, specifically *Afi-egr* and *Afi-slc4a10* show loss of expression in their respective territories (S7 Fig). Interestingly, *Afi-msp130L*, although not classified as significantly differentially expressed in the NanoString dataset but being close to log fold change −1, also showed a loss of expression in WMISH analysis. *Afi-p58b* consistently showed no change of expression quantitatively or qualitatively (S7 Fig).

**Figure 5:**
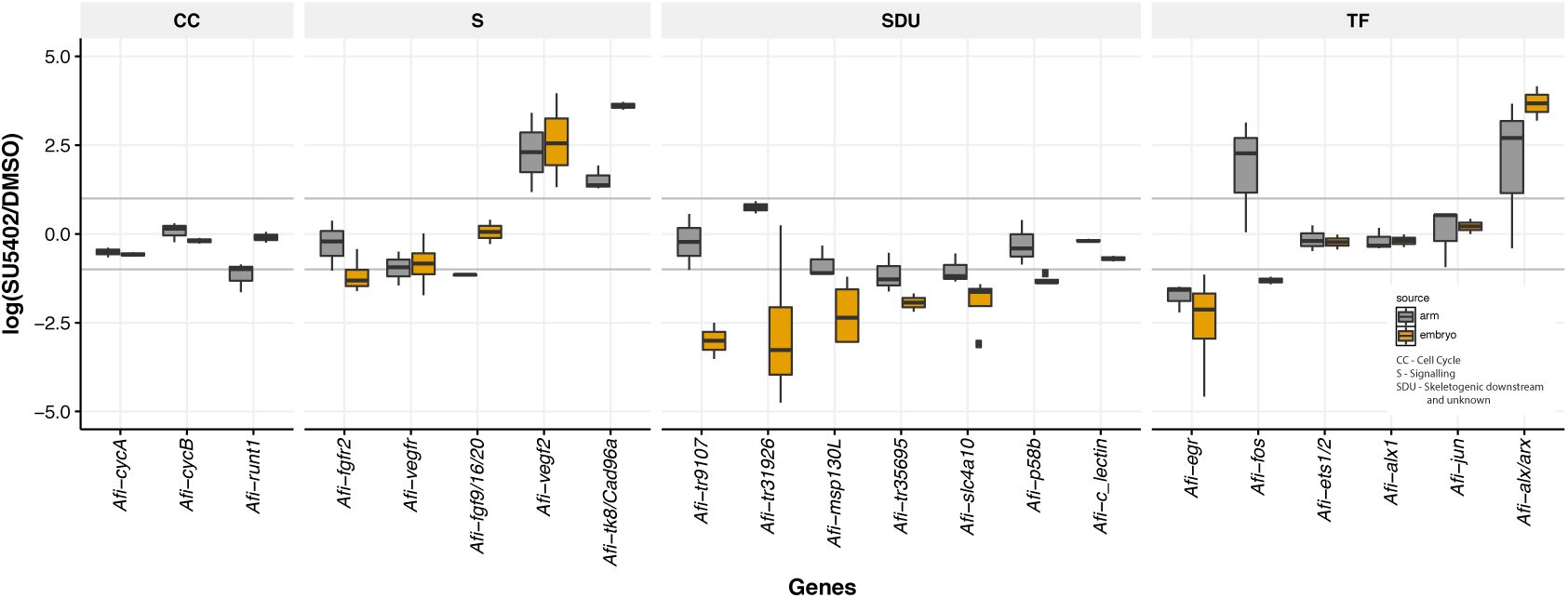
Comparison of genes affected by FGF signalling perturbation in embryos and regenerating arms of *A. filiformis*. Genes downregulated and upregulated in the same way in response to SU5402 treatment in embryos and regenerates. Threshold set is ±1 log2(SU5402/DMSO) corresponding to 2-folds of difference.

Since many candidates were found to be close to a fold change of ±1, we additionally assessed statistical significance using the Student’s t-test. We found 23 differentially expressed genes of which 7 were shared between the threshold and t-test. Due to the small overlap, not even 50%, we compared the distributions of standard deviations between arms and embryos. We found a higher dispersion in the samples collected from arms than in the samples from embryos (S8 Fig). A possible explanation for such a high variance may be that our arm samples are more heterogeneous and also contain only a small proportion of skeletogenic cells, thus increasing the noise-to-signal ratio and making it more difficult to find affected genes using standard quantitative approaches.

### Molecular effects downstream of FGF signalling in embryogenesis and regeneration

To address whether the molecular effects of FGF signalling on skeleton development are similar between embryonic development and arm regeneration, we quantitatively compared the expression of various genes in the two processes. Of the 123 genes quantified using Nanostring technology we found 24 in arm and 15 in embryo to be expressed below background (<20 counts comparable to internal negative control of the Nanostring). Using the threshold of log2(SU5402/DMSO) ±1 for a better comparison, we find that overall 73 differentially expressed transcripts show the same trend of expression between arms and embryos, with 59 downregulated (S9A Fig) and 14 upregulated (S9B Fig). 22 genes show a different trend of expression (S9C Fig). Interestingly, *Afi-msp130L* is not part of the overlapping genes in the quantitative dataset and clearly showed no expression in SU5402 treated arms or embryos by *in situ* hybridization (Fig 4, 5 and S7 Fig), suggesting that our approach may be too stringent to detect all downregulated genes, especially in the more heterogeneous context of arm regeneration. Notably, we didn’t observe any expression changes in cyclin genes (e.g. *cycA, cycD*) in agreement with the EdU analysis in the regenerates treated with SU5402 (Fig 5 and S4 Fig), in which cell proliferation is not affected. Additionally, three transcripts, homologs to the uncharacterised sea urchin tyrosine kinase *Afi-tk8/Cad96a, Afi-vegf2* and the homeodomain transcription factor *Afi-alx/arx*, are all upregulated in SU5402-treated embryos and adult regenerates (Fig 5). Interestingly, few genes differentially affected by SU5402 relative to controls in the adult regenerating arms, but not in the embryos (Fig 5), are stem cell-related transcription factors such as *Afi*-*runt1* and *Afi*-*fos*, and signalling genes belonging to other pathways (such as *Afi*-*serrate*). Importantly, upstream skeletogenic specification transcription factors (such as *Afi*-*alx1, Afi-ets1/2* and *Afi-jun*), as well as the minority of downstream skeletogenic genes (*Afi*-α*coll* and *Afi*-c*-lectin*) are unaffected by FGF signalling inhibition in both embryos and adults (Fig 5 and S9 Fig). Notably, although the signalling genes *Afi-fgf9/16/20* and *Afi-fgfr2* are expressed in comparable cell types between regeneration and development (see Fig 1), we find *Afi-fgfr2* to be downregulated only in the embryo and *Afi-fgf9/16/20* to be downregulated only in the arm, suggesting differences in regulatory processes activating the FGF signalling components during the two life stages. Finally, in both embryos and regenerating arms, FGF signalling inhibition affects the expression of VEGF signalling genes (upregulated ligand *Afi-vegf2* and mildly downregulated receptor *Afi-vegfr*), highlighting again a potential mechanism of cross-talk between the two signalling pathways. Although differences exist between development and regeneration, likely due to the anatomical differences of these two structures, we observe a high degree of similarity at the level of skeletogenic genes, suggesting similar molecular mechanisms downstream of FGF signalling.

### A new set of SM specific genes are found to be affected by FGF signalling inhibition

Impairing FGF signalling during development and regeneration severely affects the development of the skeleton in *A. filiformis*. The data in the previous sections show that a large portion of known skeletogenic genes (such as *p58a*, *kirrell*, *msp130L*) require FGF signalling to be expressed in SM cells, therefore the differential transcriptome analysis conducted on embryos treated with SU5402 can be used to identify novel downstream genes involved in the development of skeleton in *A. filiformis*. Indeed, among the 140 downregulated genes, many (27 genes) do not have a clear homolog in the sea urchin genome, used as reference for annotation. BLAST analysis, shows that a handful of these transcripts have similarities with hemichordates or cnidarian genes (S3 Table), 11 of them were included in the Nanostring codeset and analysed for their expression and response to FGF signalling inhibition in embryos and regenerating arms. Nine of these new *A. filiformis* genes show a similar response to SU5402 exposure in both the embryo and regenerating arms (S9A Fig). Bioinformatic analysis on five of these novel genes revealed that *Afi-tr31926* and *Afi-tr35695* are unique to brittle stars (also found in *Ophiocoma wendtii*) with no similarity to any other sequences within analysed echinoderms (BLAST using Echinobase/EchinoDB databases) or in other organisms (NCBI non-redundant database) (S4 Table). Protein structure prediction using PredictProtein (61) and analysis of conserved domains using CDART and PFAM databases revealed that these genes are likely to be secreted (presence of a signal peptide) and one of those (*Afi-tr35695*) is predicted to have calcium ion binding activity, which would be consistent with its putative role in the formation of a calcium carbonate skeleton (S4 Table). The spatial expression using ISH (S10 Fig) show that *Afi-tr31926, Afi-tr35695* are expressed in the skeletogenic mesoderm in both the embryo and in the regenerating arm, either in early stages, late stages or both. *Afi-tr9107*, on the other hand, is expressed in the ectoderm in a pattern that is reminiscent of the expression of the signalling ligands *Afi-fgf9/16/20* and *Afi-vegf3* in the ectoderm of the embryo at the boundary with the endoderm (Fig 4), adjacent to where the skeleton is deposited. During regeneration this transcript is expressed in vertebrae and spines of late regenerating adult arms (S10 Fig).

Interestingly, in our analysis we also found two new genes, which have not been previously described to have expression in SM cells in sea urchin. One is the transcription factor *Afi-rreb1*, not consistently downregulated in different biological replicas, and the gene *Afi-cara7la* (Fig 4C) consistently downregulated in SU5402 treated embryos. Both are specifically expressed in the skeletogenic territory in both embryos and regenerating arms and constitute additional novel skeletogenic genes identified in this study.

Altogether, these data identify new genes downstream of FGF signalling, and similarities in the molecular network driving skeletogenesis between embryonic development and adult arm regeneration (Fig 6), suggesting that they are functionally equivalent.

**Figure 6:**
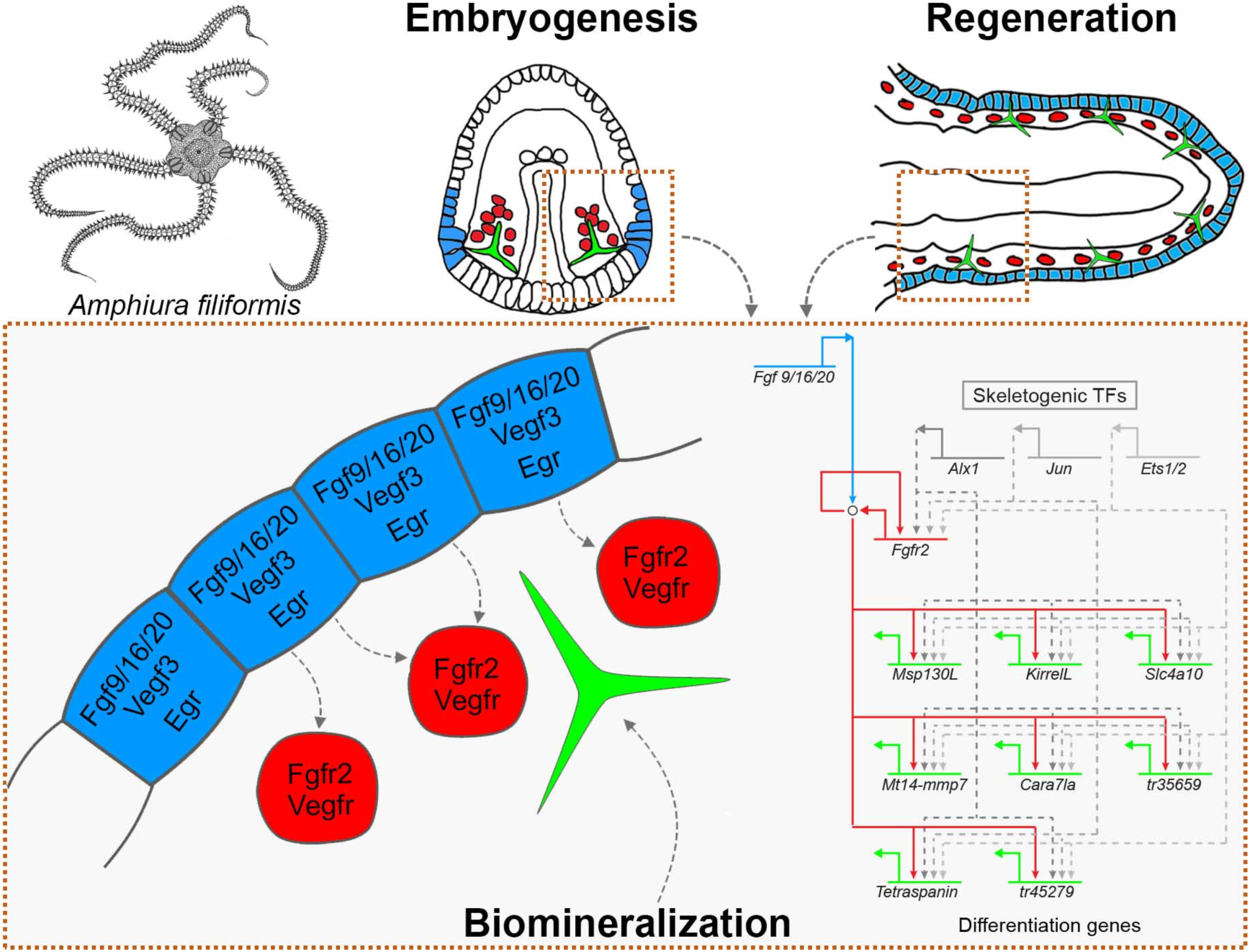
Summary of the role of FGF signalling in biomineralization and driving the skeletogenic differentiation gene battery in embryos and adult regenerating arms of *A. filiformis*. The schematic diagram represents the signalling occurring in mesenchymal and ectodermal cells. The provisional gene regulatory network shows implied connections (literature) represented by dashed lines and confirmed connections (this study) by solid lines between FGF signalling genes and biomineralization genes in the skeletogenic GRN of the brittle star

## Discussion

### FGF signalling is required for skeleton formation in the brittle star and activates a cassette of biomineralization genes

In this work we show that both brittle star skeletal development and adult regeneration rely heavily on the presence of FGF signalling. The evidence for this is as follows: 1) the expression pattern of *FGF* and *VEGF* ligands and receptors during development and regeneration allows for the ectodermal-mesodermal tissue interaction, which has been shown to be crucial for skeletogenesis in sea urchin embryos (31–34); 2) perturbation of this pathway using the universal pharmacological agent SU5402 resulted in complete inhibition of skeletal spicule formation in both adult arms and embryos; and 3) FGF signalling inhibition specifically downregulated the expression of genes involved in biomineralization. Similarly to what was suggested for sea urchins (31), we show that the role of FGF signalling during skeletogenesis in the brittle star appears to be confined to downstream differentiation of skeletogenic cells, as upstream specification transcription factors (e.g. *Afi-alx1, Afi-ets1/2*) are unaffected. Proteomic studies have revealed hundreds of proteins associated with both the sea urchin and brittle star skeletal matrices (62,63). Interestingly, FGF signalling perturbation specifically downregulated only a subset of those skeletogenic differentiation genes, while having no effect on others (e.g. *Afi-p19, Afi-c-lectin*). Nevertheless, this subset of downregulated genes is essential for skeleton formation as their collective downregulation results in the failure of the last checkpoint in the skeletogenesis network – deposition of the biomineralized skeleton by mesenchymal cells. Those include genes belonging to the carbonic anhydrase gene family (e.g. *cara7la)*, implicated in calcium carbonate deposition in various organisms including sea urchins (64,65) and molluscs (66), solute carrier proteins like *slc4a10*, and mesenchymal surface glycoproteins like *msp130* (67).

### Functional conservation of FGF signalling in embryonic and regenerative skeletogenesis

A molecular conservation of genes expressed during embryonic development and regeneration was previously shown in newts (3) and chick embryos (4). However, these studies were limited to a comparison of only one or a few genes. Most recently, the transcriptomes of the embryo and regenerating stages of the sea anemone, *Nematostella vectensis*, have provided the first large-scale resource for comparing those two processes (7). It is thus of great interest to compare specific aspects of development between embryogenesis and regeneration. Our previous work has already shown that the morphology and molecular signature of skeletogenic cells is highly similar between the embryo and regenerating adult arm of *A. filiformis* (35,37,38). Our results showing the importance of FGF signalling in skeleton development and regeneration in *A. filiformis* reveal additional functional similarities between skeletogenesis at both stages of the brittle star life-cycle. The underlying molecular network downstream of FGF signalling is highly conserved between regeneration and development, with 15 genes being specifically downregulated in both cases e.g. the transcription factor *Afi-egr* or the biomineralization genes *Afi-kirrelL, Afi-msp130L* and *Afi-slc4a10*. Additionally, a previously uncharacterized tyrosine kinase receptor (*Afi-TK8/Cad96a*), *Afi-vegf2* and *Afi-alx/arx* are all upregulated in both embryos and adult arms. Taken together, our data provides support for the hypothesis of regeneration re-capitulating development, at least at the level of cell differentiation, and provides the first large-scale comparison of the molecular networks driving development and regeneration of the same species. It remains to be found whether the initiating signals upstream of this signalling pathway are conserved between embryonic development and adult regeneration.

### Cross-talk between FGF and VEGF signalling regulatory networks

It has been previously suggested that the FGF and VEGF signalling pathways may function synergistically, whereby the downregulation of either of the ligands can affect the expression of the other pathway components (32,68). For instance, specifically in sea urchins, downregulation of *fgfa* results in upregulation of *vegf3* expression. Our large-scale analysis of downstream targets of the FGF pathway provides insights into the mechanisms of its transcriptional regulation. Our results show that the inhibition of FGF signalling impinges on the expression of VEGF pathway genes (upregulation of ligand *Afi-vegf2* and downregulation of the receptor *Afi-vegfr10*, expressed in skeletogenic cells). Additionally, we found that treatment with SU5402 upregulates another tyrosine kinase receptor gene, *Afi-tk8/Cad96a*, closely related to *Drosophila Cad96a*. Altogether, this data underlines the difficulty with dissecting the roles of signalling pathways, which may be tightly linked to inter-regulatory loops, suggesting the presence of a signalling network in which ligands and receptors are under the control of other signalling pathways.

### Evolution of FGF signalling and skeleton formation in echinoderms

The evolution of the FGF gene family has been shown to involve extensive gene duplication and gene loss, often lineage-specific (69), resulting in complex and variable distribution of FGF genes amongst the metazoans. Only one FGF ligand has been described in sea urchins, belonging to the FGF9/16/20 subfamily, whereas hemichordates have five ligands (70), some of which result from specific duplications within the ambulacraria. In vertebrates, major duplications of the gene family occurred resulting in 19 FGFs in chicken and over 22 FGFs in mammals (69,71). There are far fewer receptors of the FGF signalling pathway with only four functional FGFR genes in vertebrates (72), two genes in sea urchins (73) and two in *Drosophila* (72). We found that the *A. filiformis* transcriptome has two clear FGF ligands, one belonging to FGF9/16/20, the other to FGF8/17/18 so far not described in sea urchin. Our results are consistent with previous studies suggesting extensive duplications in the vertebrate lineage (70) during evolution of the FGF gene family.

In sea urchin embryos, FGF signalling components are expressed in a complimentary pattern, whereby the *fgfr2* receptor is specifically expressed by the skeletogenic mesoderm cells and the *fgfa* ligand is expressed in overlying ectoderm flanking those cells (31,32). Recent work showed that this pattern of expression is also observed for the VEGF signalling genes in both sea urchin (32–34) and brittle stars embryos, as well as in sea urchin and sea star juveniles (74,75). It has been suggested that the heterochronic activation of this pathway in sea urchin and brittle star embryos lead to the co-option of the adult skeleton into the larva (74,76), as sea star embryos do not have those genes expressed at the embryonic stage and have no larval skeleton (74). Our results show that both VEGF and FGF genes are expressed in a strikingly similar pattern in embryos and adult regenerating arms of *A. filiformis*, suggesting that the interaction of the skeletogenic cells with the ectoderm, mediated by those signalling pathways, may be a conserved feature for adult echinoderms, and has in fact been co-opted in the embryos of sea urchins and brittle stars to form a larval skeleton. It is important to notice that our data suggest in brittle stars FGF signalling plays a more prominent role in skeletogenesis than VEGF signalling, which is the opposite case for sea urchins (32). Furthermore, the transcriptional regulation downstream of FGF signalling appears to be significantly different in brittle stars and sea urchins. We found the following lines of evidence for this: 1) approximately 30% of genes identified in our differential screen did not have sea urchins homologs (e.g. *tr31926* and *tr35695*); 2) other genes with homologs are not specifically expressed in the skeletogenic lineage in the sea urchin (e.g. *Afi-rreb1;* (77)). Recent work showed that despite a striking similarity in the morphology and development of the larval skeleton in sea urchins and brittle stars, the dynamics of their regulatory states are very different, suggesting alternative re-wiring of the network in the two classes (35). Together with our results showing the high degree of conservation of the brittle star embryonic and adult network downstream of FGF signalling, we can hypothesize that the embryonic program for skeletogenesis could have been independently co-opted in brittle stars and sea urchins. An alternative evolutionary scenario would imply a coordinated evolution of the skeletogenic program in larvae and adult. Elucidating the role of FGF signalling in adult skeletogenesis of the remaining four classes of extant echinoderms could help resolve this issue in the future.

### Evolutionary implications for skeletogenesis among deuterostomes

Skeletal regeneration is observed in other deuterostome groups: for example in cirri regeneration of amphioxus (78), and in appendage regeneration of different vertebrates (reviewed in (79)). It has even been suggested that adult bone repair and regeneration may recapitulate embryonic bone development at a molecular level (80). Comparing the skeleton developmental program between embryogenesis and regeneration can be vital to understand the evolution of the skeleton in deuterostomes. Although the skeleton of echinoderms is composed of calcium carbonate, instead of calcium phosphate, similarities of its ontogeny can be observed when compared to vertebrates. For example, in both groups of animals the trunk skeletal precursor cells are mesoderm-derived, motile mesenchymal cells (35,53,81,82), already suggesting some conserved features of skeletogenesis in deuterostomes. Gene expression can also aid in understanding the extent of potential similarities. The key regulators of the sea urchin (and likely brittle star) skeletogenic GRN include transcription factors *alx1*, *ets1/2* and *erg* (35,37,39,83–85). Members of the Cart/Alx3/Alx4 group of transcription factors are also involved in skeletal development in vertebrates. They are expressed in embryonic lateral plate mesoderm, limb buds, cartilage and ectomesenchyme, and deletions of these genes result in cranial and vertebral malformations (40,41,86). ETS family transcription factors (including homologs of *ets1/2* and *erg*) have also been implicated in vertebrate skeletogenesis (87–92). FGF signalling has a highly conserved role in skeletogenesis in deuterostomes, as demonstrated in sea urchins (31) and lampreys (93), as well as chickens (94) and mice (95,96).

Overall, in terms of the downstream biomineralization genes, it seems that the network has diverged significantly between echinoderms and vertebrates. Most of the biomineralization genes identified in sea urchins and brittle stars do not have apparent homologues in vertebrates or other invertebrate deuterostomes (35,63,65). Interestingly, the recent genome of the brachiopod *Lingula anatine*, which like distantly related vertebrates forms its shell using calcium phosphate, also reveals a unique expansion of a set of biomineralization genes (for example chitin synthases) different from duplication events which gave rise to bone formation genes in vertebrates (such as fibrillar collagens) (97). Those differences in the set of biomineralization genes used by brachiopods, echinoderms and vertebrates suggest that these animals independently evolved a core differentiation gene cassette via duplication events for building their calcium-based skeletons. Nevertheless, the initiation cascade, including the ancient signalling pathways (e.g. FGF, BMP) and transcription factors, appears to play a conserved role in these divergent animals (65,97–100). Together with these studies, our work presents further evidence for an evolutionary conserved regulatory apparatus driving the activation of biomineralization genes.

## Conclusions

In this study, we show for the first time a comparison of the role of FGF signalling in the embryonic development and adult regeneration of the same species, the brittle star *Amphiura filiformis*. We characterized the expression of FGF and VEGF signalling pathway ligands and receptors during both embryonic development and adult arm regeneration. Using the pharmacological inhibitor SU5402 we showed that perturbation of FGF signalling interferes with skeleton formation during both developmental processes. Our transcriptome-wide analysis of the effects of FGF signalling inhibition in brittle star embryos revealed a global view of the downstream targets of this pathway, including well-studied genes as well as novel brittle star skeletogenic genes. Finally, our comparative analysis of these FGF targets between embryos and adult regenerating arms strongly supports a high degree of conservation of the downstream molecular network underlying skeletogenesis. Identification and comparison of the upstream signals initiating the skeletogenic gene regulatory network in embryos and adults will elucidate whether regeneration re-capitulates development, as well as contribute to our understanding of the evolution of skeletogenesis within both echinoderms and deuterostomes more broadly.

## Materials and Methods

### Adult animal maintenance and handling

Adult animals of *Amphiura filiformis* were collected during their reproductive season (July-August) for embryo cultures and throughout the year for adult specimens in the Gullmarsfjord, Sweden in the proximity of the Sven Lovén Centre for Marine Sciences. Animals were maintained in the laboratory in London as described previously (37). Regenerating arm samples were obtained as described (38) while amputated arm explants were obtained as described in Burns et al., 2012. *A. filiformis* embryo culture was set up as previously described (101). Treated and untreated embryos were collected at required stages for WMISH, RNA extraction and RNAseq as previously described (35,36).

### Whole mount in situ hybridization

The protocol for WMISH for embryos and adult regenerating arms of *A. filiformis* was identical except for the hybridization temperature as outlined below. The samples were first re-hydrated with graded ethanol washes (70%, 50% and 30%) and washed three times in 1x MA Buffer with Tween (MABT; 0.1M Maleic Acid pH 7.5, 0.15M NaCl, 0.1% Tween-20) and pre-hybridized in hybridization buffer (50% deionized formamide, 10% PEG, 0.05M NaCl, 0.1% Tween-20, 0.005M EDTA, 0.02M Tris ph 7.5, 0.1mg/ml yeast tRNA, 1x Denhart’s solution, DEPC-treated water) for 1h at 50°C (regenerating arms) or 55°C (embryos). Next, the samples were put in HB containing 0.2ng/*μ* l antisense probe for 3-7 days at the same temperature. Following this period of time samples were post-hybridized in fresh HB without probe for three hours, then washed once in MABT at the corresponding hybridization temperatures and once at room temperature (RT). The samples were then washed three times in 0.1x MABT, once more with 1x MABT before placing them in blocking solution (MABT, 0.5% goat serum) for 30 min. Samples were then incubated in 1:1000 anti-DIG AP (Roche) antibody solution overnight at 4°C. Next, the sample was washed 5 times in 1x MABT and 2 times in alkaline phosphatase (AP) buffer (Tris pH 9.5, MgCl_2_, NaCl, Tween-20, levamisole, milliQ water) before adding the staining solution (AP buffer, 10% DMF, 2% NBT/BCIP) for the chromogenic detection. The staining was stopped with MABT washes.

### Inhibitor treatments and phenotypic analysis

SU5402 (Calbiochem) and Axitinib (Sigma) were dissolved in DMSO for a stock concentration of 10mM and 5mM respectively. The drugs were added to embryo cultures at 18hpf at a final concentration of 10μM (SU5402) and 75nM (Axitinib), and the embryos were then allowed to develop until 27hpf. At this time-point approximately 500 treated and control (0.2% DMSO and FSW) embryos were collected for fixation for *in situ* hybridization and 500 embryos were collected for RNA extraction for QPCR and NanoString analysis. Remaining embryos were left to develop further for phenotypic assessment. Arm explants were used for testing the effects of inhibitors on regeneration and skeletogenesis. Adult *A. filiformis* arms were cut 1 cm from the disc and then left to regenerate until stage 2 (on average 5 days post-amputation). Arms were then cut again 5mm proximally to the initial amputation site to obtain explants, which were left for several hours to allow proximal wound healing. The explants were then incubated for 24h in SU5402 at a final concentration of 10μM or Axitinib at 200nM. Samples reared in FSW and 0.1% DMSO were used as controls. To determine phenotypic effects on spiculogenesis, calcein staining (Sigma) was used together with the inhibitor to label any newly forming spicules (1:50 dilution of a 1.25mg/ml stock solution). After the treatment, the arm explants were imaged for any morphological phenotype and fixed for ISH or collected in RLT (15 arms per condition) for NanoString analysis.

### Differential analysis of transcriptome data

Samples were quantified and normalized as previously described (36). Differential analysis was conducted between SU5402 and DMSO treated samples. Since only one biological replicate was used for mRNA-seq, we used it to identify potentially differentially expressed candidates and validated those using other technologies. Candidates were selected based on two criteria: (1) a user defined threshold of expression above 2 tpm and (2) a fold-change threshold of +-1.6.

Since all three methods (transcriptome quantification, NanoString and QPCR) employ technologically different quantification strategies, we assessed their technical similarity by comparing fold change values of the different methods on the same biological replicate. Consistently, all three technologies showed a similar trend in fold change (88.1%). Transcriptome and NanoString fold change values for 114 genes showed high positive correlation (~0.85) and linear regression analysis resulted in a significant positive association between the two techniques (β=1.12, 95% CI [0.97, 1.24], p***<0.001, adjusted R^2^=0.7223; S6 Fig). Interestingly, fold change values seemed generally slightly inflated in the transcriptome dataset (slope > 1). When comparing fold change values of 31 genes between QPCR and Transcriptome we found a positive correlation (~0.854) and that both techniques are positively associated (β=1.4340, 95% CI [1.10, 1.77], p***<0.001, adjusted R^2^=0.7203; S6 Fig). This is consistent with our observation comparing correlations of time-course datasets quantified using transcriptomics, NanoString and QPCR (36). Importantly, since every technology encompasses differences in their technical error, we used the β and y-intersect values of the linear regression analysis to compare biological replicates across technologies.

### Inference of phylogenetic gene trees

For phylogenetic gene trees, sequences were collected from local assemblies and publicly available datasets (41 species). To fish out genes that contained the FGF, FGFR, VEGF and VEGFR domains, we obtained HMM profiles from the PFAM database. The sequences of the 41 species were scanned against these domains and were used to generate input data for OMA (v2.2.0) (102). Hierarchical orthologous groups that contained our candidates were merged with groups that showed close blast similarity and were selected for further analysis. The merging step was necessary due to the independent divergence between chordates and echinoderms of more than 500 Mya (million years ago) and still too low taxonomic sampling. Mafft (v7) (103) was used for multiple sequence alignment, followed by several manual rounds of sequence trimming using maxAlign (v1.1) (104) or independent criteria such as retention of close sequence length to given candidate. For tree inference we used Iqtree (v.1.5.5) with LG model and 1000 fast bootstraps (105).

### Validation of differentially expressed candidates using QPCR and NanoString

To validate candidates obtained from the transcriptome analysis we performed a linear regression analysis between transcriptome vs QPCR and transcriptome vs NanoString using R. Coefficients obtained for slope and y-intercept were used to scale QPCR and NanoString samples in relation to the transcriptome. In this way, we accommodate differences in intrinsic technical errors of the various technologies.

### QPCR and NanoString nCounter analysis

QPCR analysis was performed as described previously for adult regenerating arms (37) and embryonic samples (35). Additionally, differential expression of genes was measured using the NanoString nCounter analysis system (NanoString Technologies, Seattle, WA, USA) (106). A 123-probe code set was designed based on *A. filiformis* sequences, including six different internal standard genes and a GFP probe for detecting spike-in GFP RNA (S2 Table). For each experimental sample 100ng of total RNA was used, extracted from 300 embryos and 10 regenerating arms respectively, using the RNeasy Micro Kit (Qiagen). Detected counts/100ng of total RNA were normalized first using the positive control lane normalization provided in the NanoString nCounter cartridge and then again using selected six internal standard genes (normalization factor obtained using geometric mean for each lane). For quantifying differential gene expression in perturbed samples, a Log2 fold change between controls and treated samples was calculated. The Log2(SU5402/DMSO) of ±1 (reflecting a 2-fold difference in change of level of expression) was determined to be biologically significant in correspondence with previously published work (60).

### Cell proliferation assay

Regenerates treated with the SU5402 inhibitor were tested for changes in cell proliferation. The cell proliferation assay was carried out using the Click-iT® EdU HCS Assay (Invitrogen) as described previously (38) then imaged using confocal microscopy. For each regenerate between ~100 ±10 slices were taken per Z-stack (1µm thickness). DAPI-labelled nuclei and EdU-Labelled nuclei per stack were counted automatically using the Fiji plugin TrackMate (107). Number of EdU labelled nuclei per total number of nuclei ranged from 672/3375 to 1385/5205. The proportion of nuclei labelled with EdU compared to all nuclei labelled with DAPI was calculated as a percentage. Student’s T-test was used and showed no significant difference between control (DMSO) and SU5402-treated samples (T-value = 0.261; p>0.25).

## Acknowledgments

We would like to thank the staff at the Sven Lovén Centre for Marine Sciences in Kristineberg, especially Sam Dupont, for assistance during animal and sample collection. We thank Olga Ortega-Martinez for sharing the regenerating explant collection method. We would like to thank Jeffrey Thompson, Johannes Girstmair and Maria Kotini for helpful comments on the manuscript.

## Author contributions

AC, DVD and PO conceived and designed the experiments. AC, DVD and LC carried out experiments. AC, DVD and PO analysed the data. AC, DVD and PO wrote the manuscript.

