## Supplemental figures for "FGF signalling plays similar roles in development and regeneration of the skeleton in the brittle star *Amphiura filiformis*"

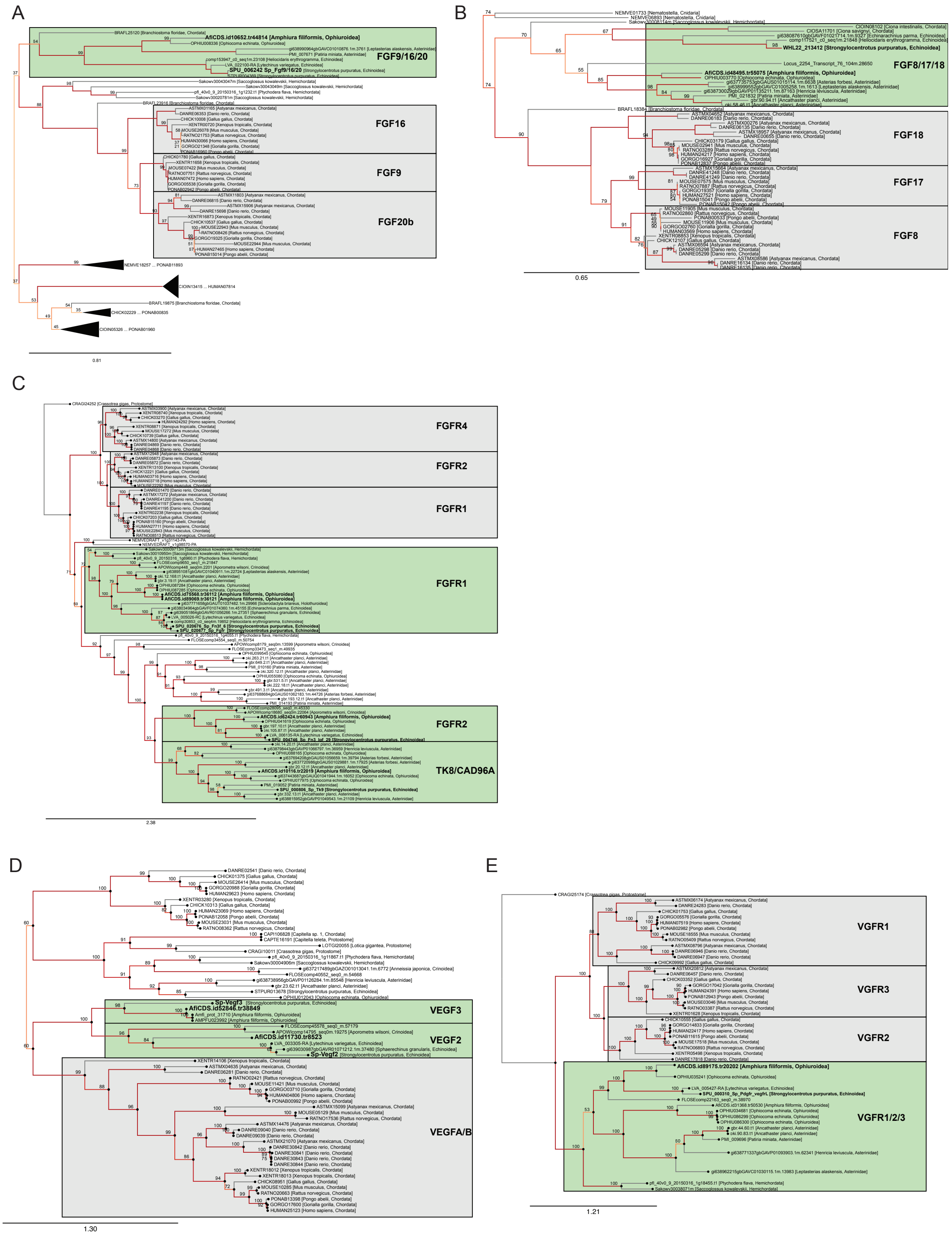


**Figure S1: Gene trees for key FGF/FGFR and VEGF/VEGFR genes**. A) AfiFgf9/16/20 is orthologous to sea urchin Sp-Fgf9/16/20 gene and form a sister group to human FGF9, FGF16 and FGF20 genes. B) Afi-Fgf8/17/18 is orthologous to unclassified sea urchin gene and form a sister group to human FGF8, FGF17 and FGF18 genes. C) Afi-Vegfr is orthologous to sea urchin Sp-Vegfr gene and is homolog to 3 independently duplicated chordate Vegfr genes (VEGFR1, VEGFR2 and VEGFR3). D) Afi-Fgfr1 and Afi-Fgfr2 are orthologs to sea urchin show independent duplication from a common fgfr gene in metazoans. E) Afi-Vegf2 and Afi-Vegf3 are clear orthologs to their sea urchin counterparts and are descendants from an ancestral Vegf gene that got independently duplicated in chordates. On nodes are fast bootstrap values (101).

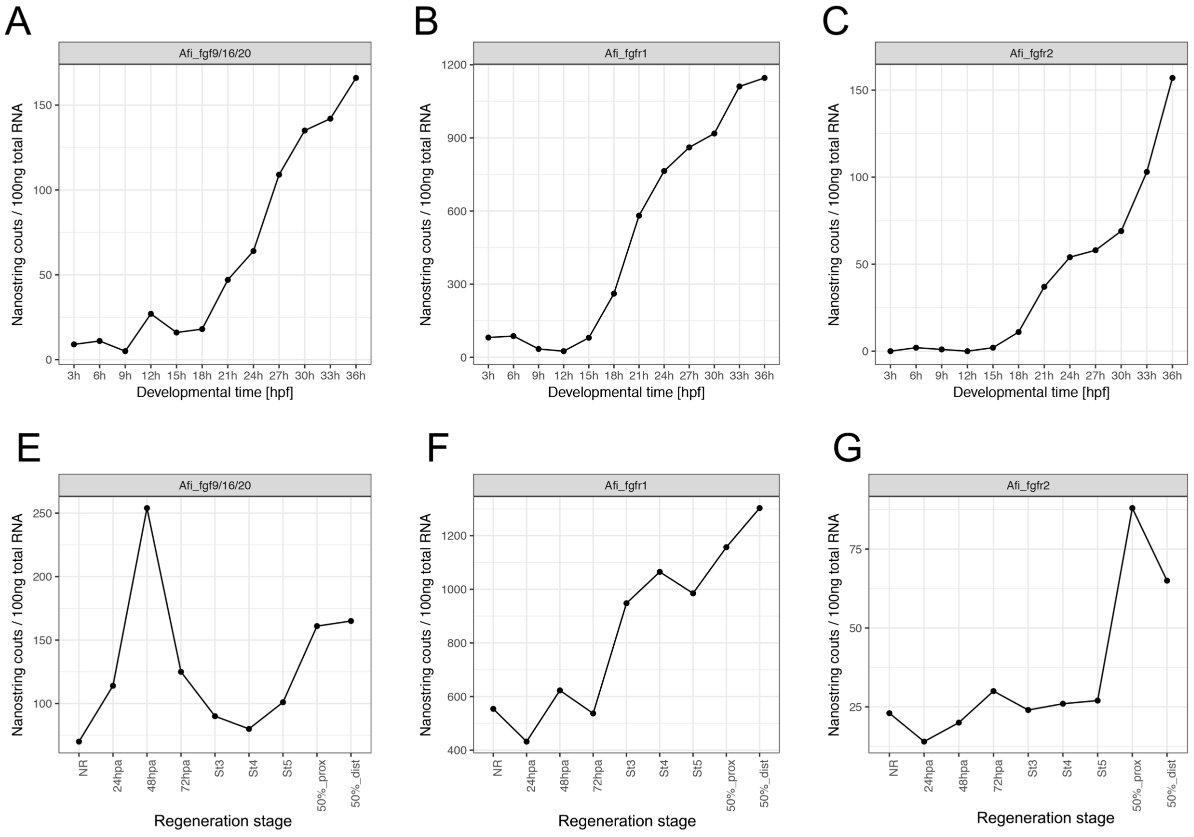


**Figure S2: Timecourses of FGF component genes during embryonic development and arm regeneration stages in *A. filiformis***. A-C) Expression of A) *Afi-fgf9/16/20*, B) *Afi-fgfr1* and C) *Afi-fgfr2* in embryos. D-E) Expression of D) *Afi-fgf9/16/20*, E) *Afi-fgfr1* and F) *Afi-fgfr2* in adult non-regenerating and regenerating arms at different stages. Transcript abundance is represented as Nanostring counts/100ng of Total RNA. Hpf – hours post fertilization, hpa – hours post amputation, prox – proximal, dist – distal, NR – non-regenerating.

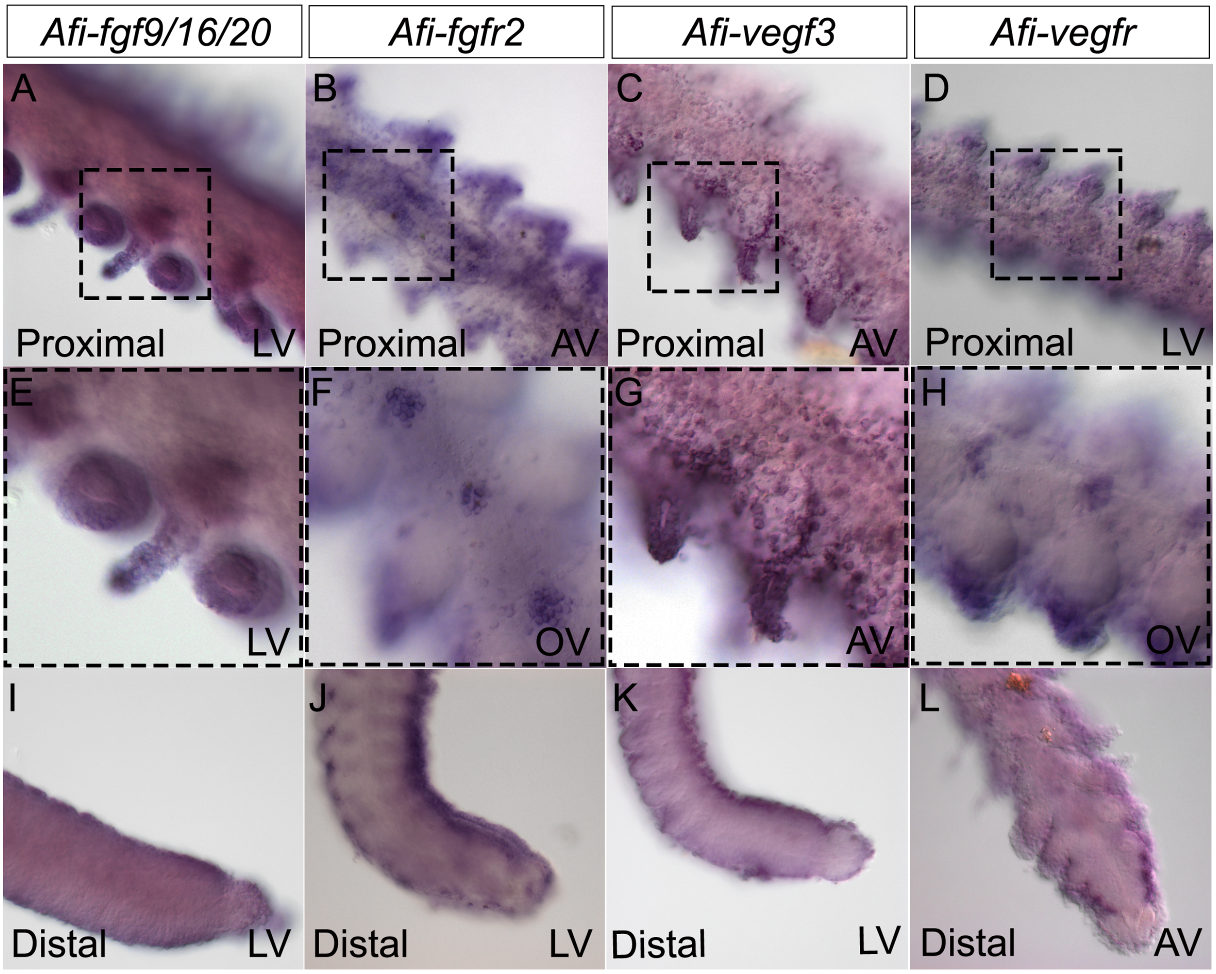


**Figure S3: Expression of FGF and VEGF genes at late stages of arm regeneration in *A. filiformis***. Av –aboral view, OV –oral view, LV – lateral view.

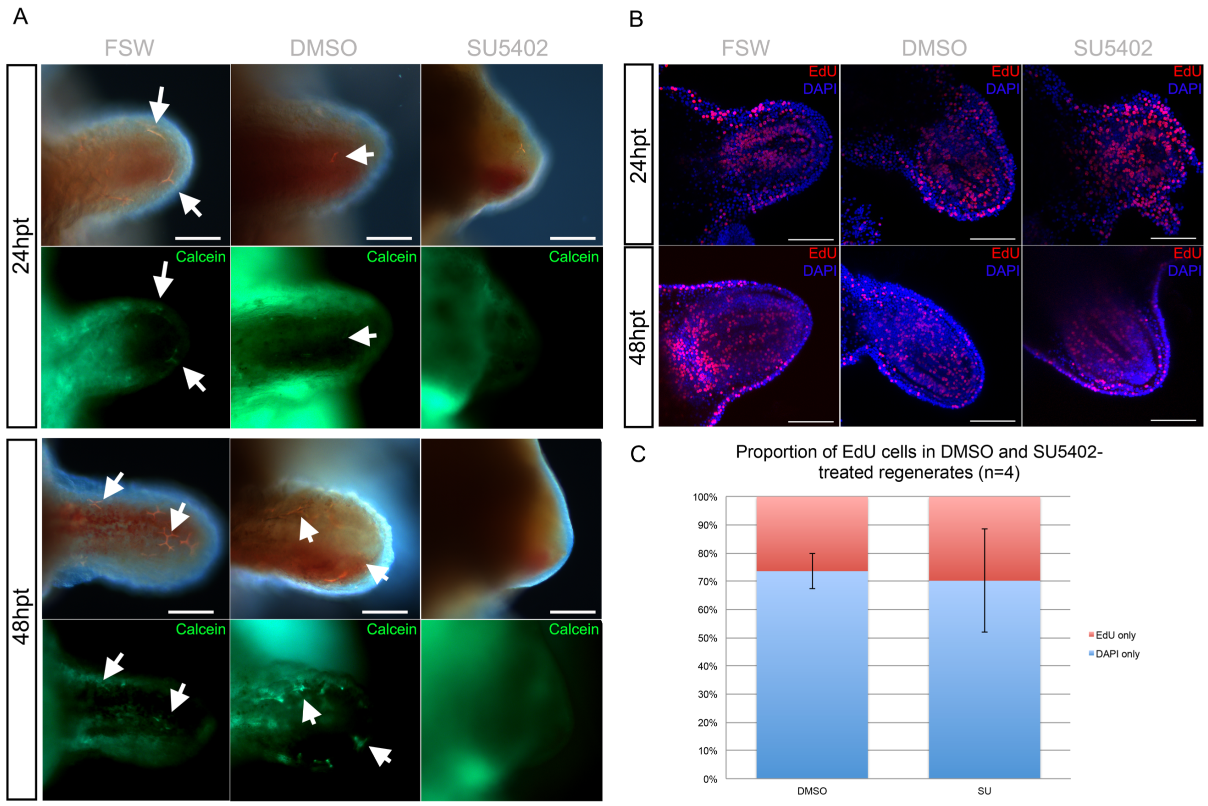


**Figure S4: FGF signalling perturbation interferes with arm regeneration in *A. filiformis* but not by reducing cell proliferation**. A) Phenotypic analysis of regenerating arm explants in control (FSW and DMSO) and SU5402 conditions at 24 hours post treatment (hpt) and 48hpt shows that skeletogenic spicules do not form and the arm seizes to regenerate further. B) Confocal images of an EdU cell proliferation assay on control and treated regenerates shows no changes in the number of EdU labelled nuclei in SU5402-treated explants both at 24hpt and 48hpt. C) Quantification of the results in B showing no significant decrease in the proportion of EdU-labelled nuclei relative to all the nuclei counted.

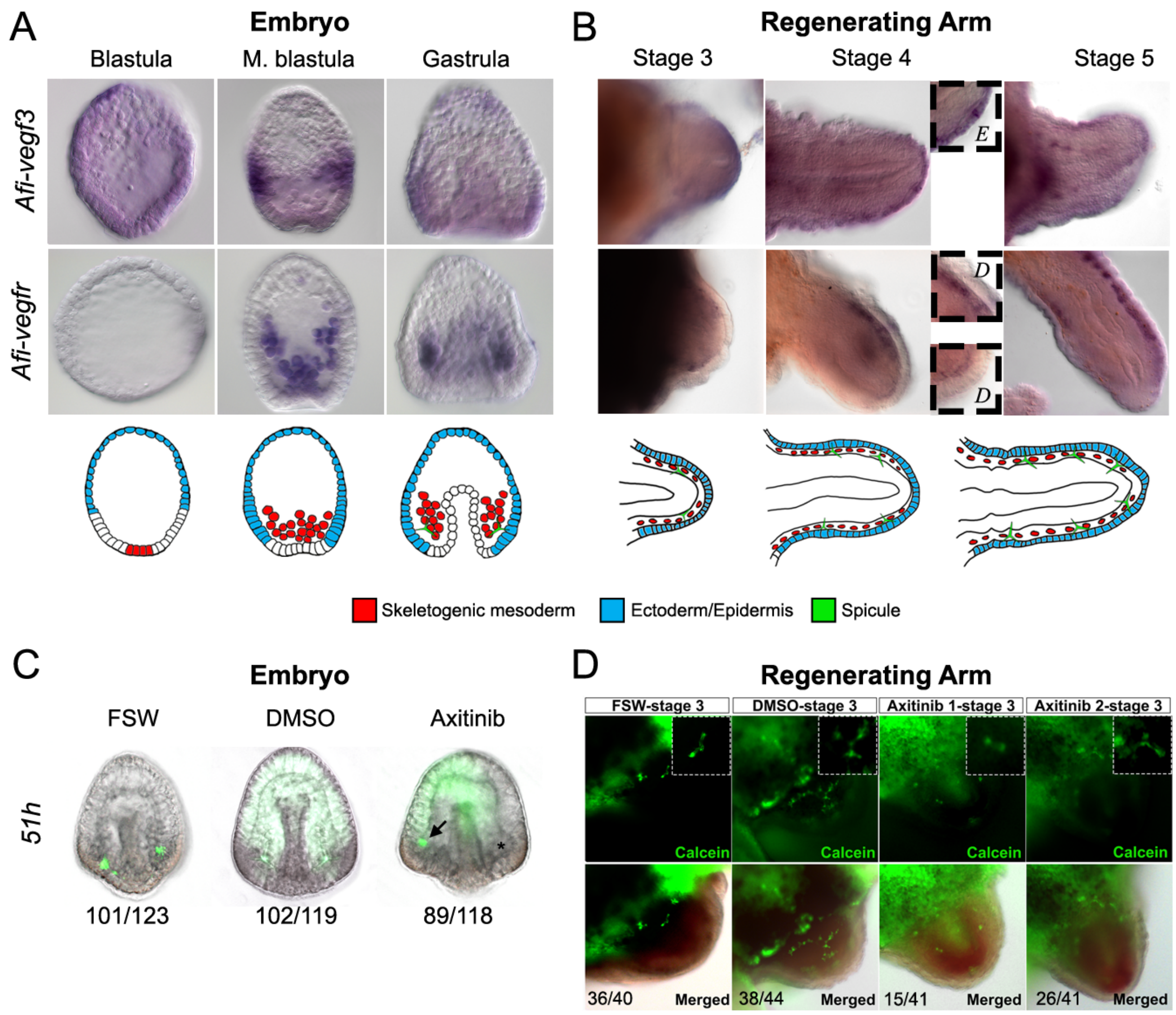


**Figure S5: Expression of VEGF signalling components and axitinib treatment in embryos and early regenerating arm stages of *A. filiformis*.** A) Top: WMISH on embryos at blastula, mesenchyme blastula and gastrula stages of development showing the expression of *Afi-vegf3* and *Afi-vegfr*. Bottom: schematic diagram of major relevant cellular domains. B) Top: WMISH on regenerates at stages 3, 4 and 5 showing the expression of *Afi-vegf3* and *Afi-vegfr*. Insets show detail of expression patterns. Bottom: Schematic diagram of major relevant cellular domains in regenerates. C) Phenotypic analysis of axitinib-treated embryos and controls at 51 hpf shows that perturbation of VEGF signalling results in embryos with one skeletal spicule forming. Numbers at the bottom show counts for embryos observed with the represented phenotype/total embryos counted. D) Phenotypic analysis of axitinib-treated regenerates and controls at 24 hours post treatment (stage 3) show that perturbation of VEGF signalling either results in normal or slightly reduced skeletal spicules. Numbers at the bottom show counts for explants observed with the represented phenotype/total explants counted. Insets show magnification of skeletal phenotypes.

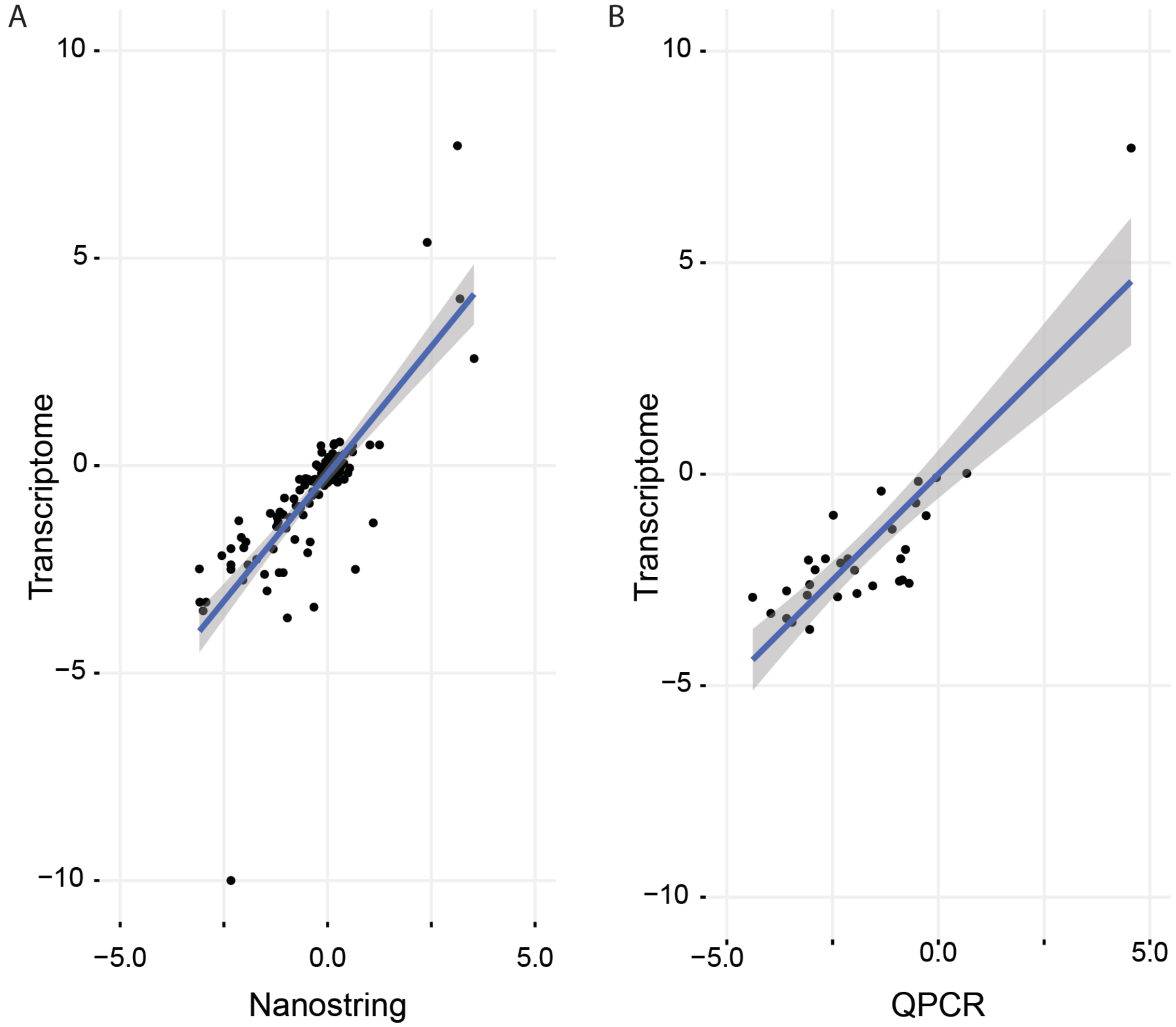


**Fig S6:** A) Correlation between Transcriptome and Nanostring quantification strategies in embryos of *A. filiformis*. B) Correlation of Transcriptome and QPCR quantification strategies in embryos of *A. filiformis*.

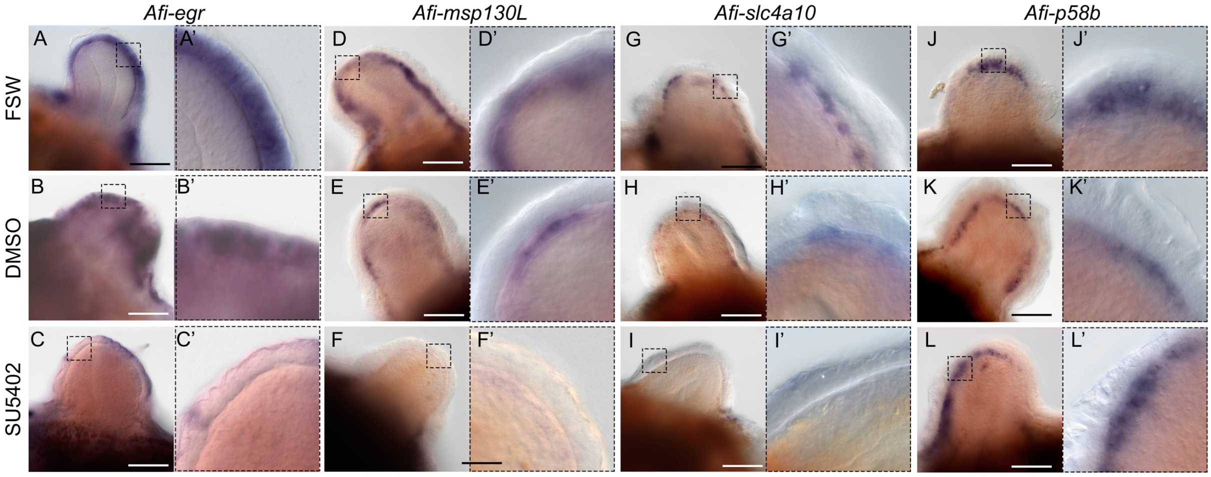


**Figure S7: Spatial downregulation of selected genes in regenerating arm samples treated with SU5402 compared to controls**. A-C) WMISH on regenerating arms shows normal expression of *Afi-egr* in epidermis is downregulated in SU5402-treated samples. A'-C') Same images at higher magnification. D-F) WMISH on regenerating arms shows normal expression of *Afi-msp130L* in the skeletogenic dermal layer is downregulated in SU5402-treated samples. D'-F') Same images at higher magnification. G-I) WMISH on regenerating arms shows normal expression of *Afi-slc4a10* in the skeletogenic dermal layer is downregulated in SU5402-treated samples. G'-I') Same images at higher magnification. J-L) WMISH on regenerating arms shows normal expression of control gene *Afi-p58b* in the skeletogenic dermal layer is maintained SU5402-treated samples. J'-L') Same images at higher magnification. Scale bars: 100µm.

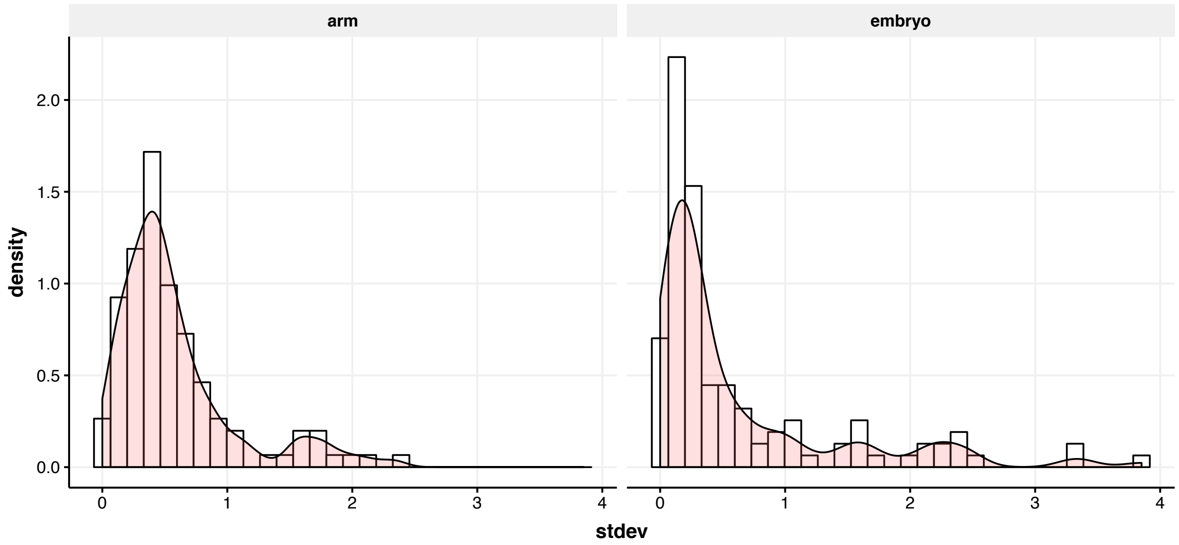


**Figure S8:** The distributions of standard deviations of differential expression in SU5402 samples between arms and embryos.

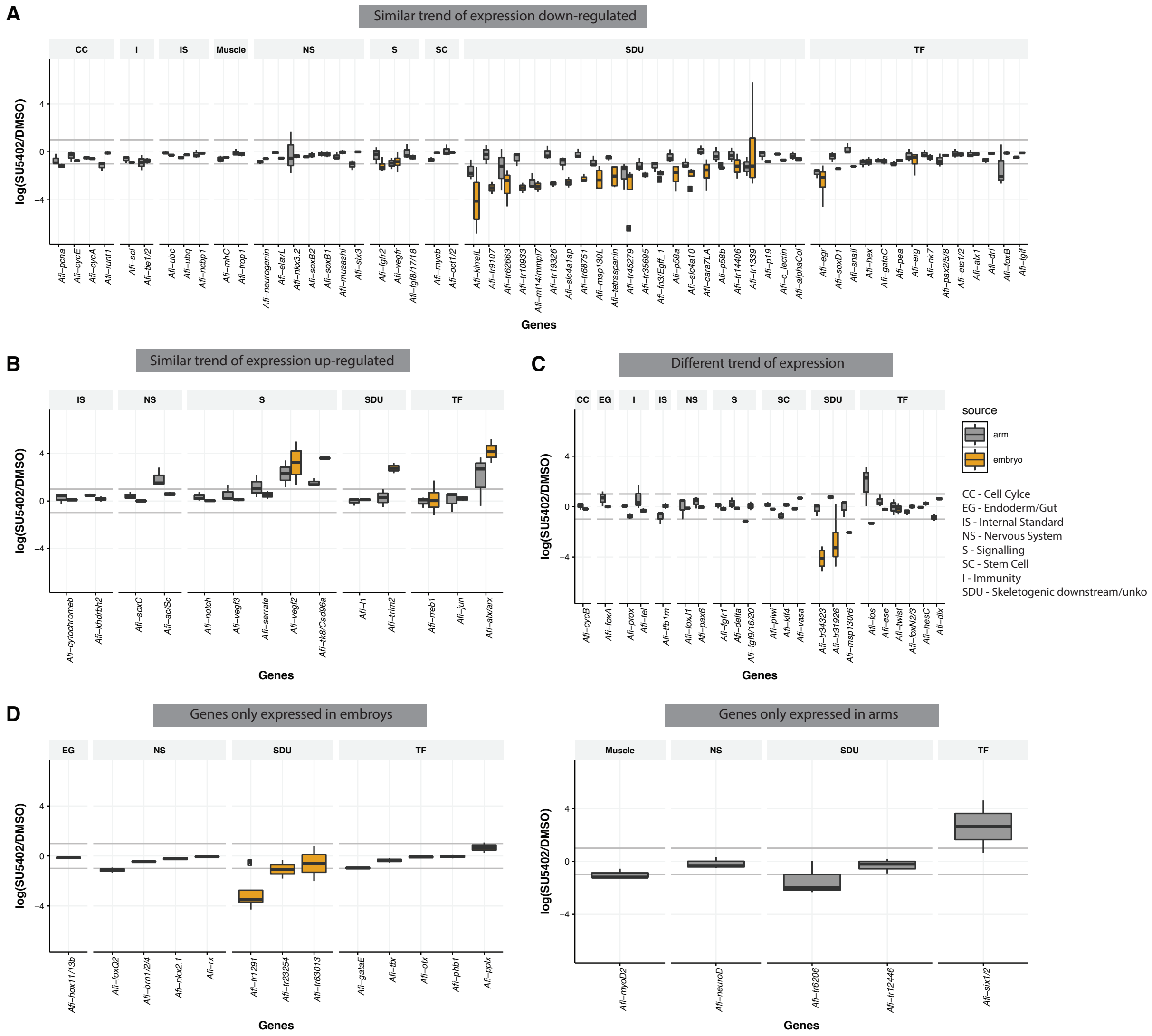


**Figure S9: Summary of the comparison of genes affected by SU5402 treatment in embryos and regenerating arms of the brittle star**. Genes showing similar trends between embryos in regenerates that are A) downregulated or B) upregulated. C) Genes showing different trends in expression between embryos and regenerates. Genes expressed above background levels exclusively during D) development or E) regeneration.

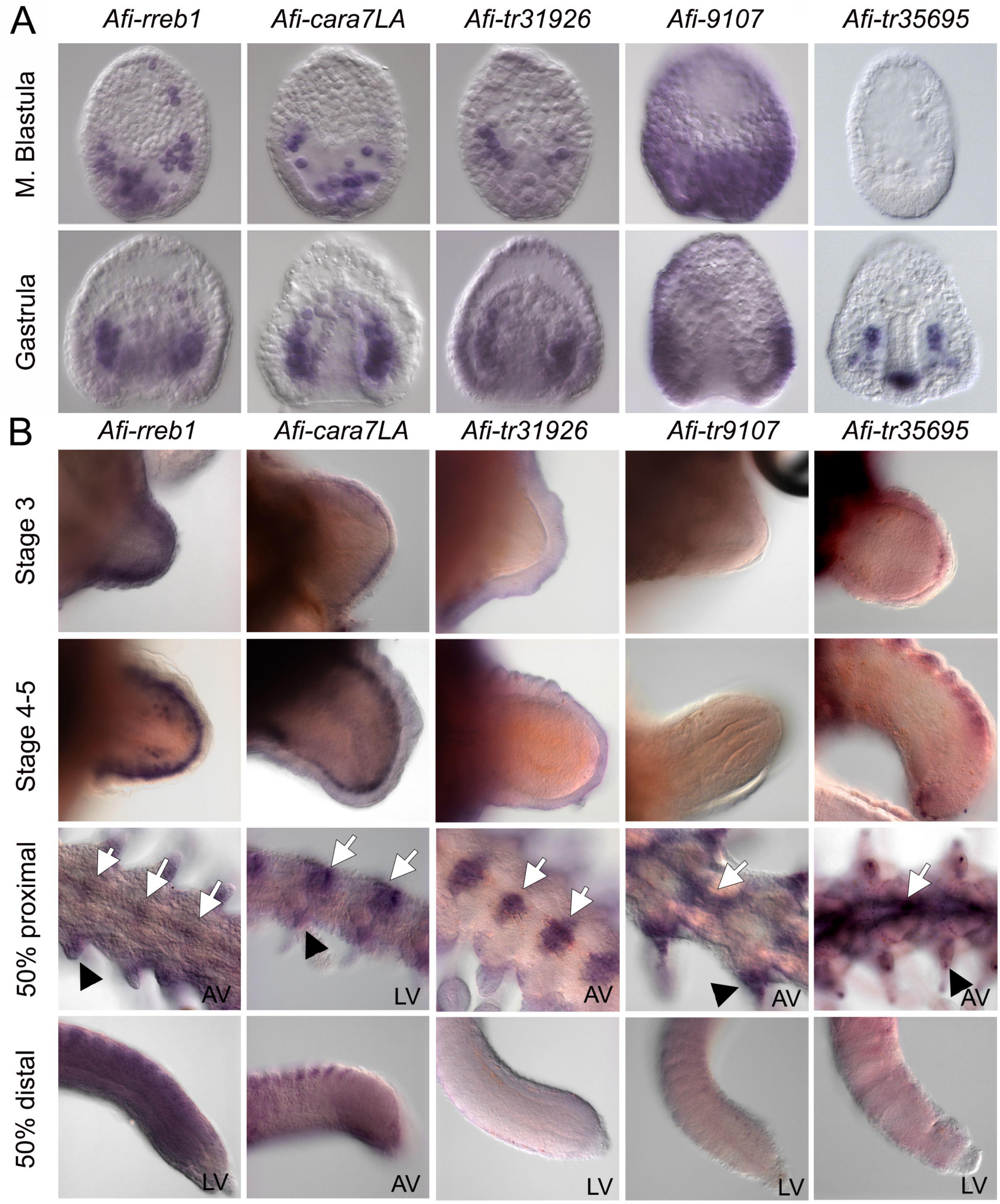


**Figure S10: Novel genes in skeletogenesis.** Expression pattern of *Afi-rreb1, Afi-cara7La, Afi-tr31926, Afi-tr9107,* and *Afi-tr35695* at mesenchyme blastula and gastrula stages of embryogenesis and early and late stages of adult arm regeneration in the brittle star. White arrows – expression in vertebrae, Black arrowheads – expression in spines. LV – lateral view, AV –aboral view.

**Movie S1: Control and SU5402-treated regenerating arm explants are alive and motile after 48h of treatment.**
