## Supplemental Table S2 for "FGF signalling plays similar roles in development and regeneration of the skeleton in the brittle star *Amphiura filiformis*"

| **Gene name** | **NCBI BLAST (NR)^1^** | **E-value** | ***EchinoDB BLAST**** | ***E-value*** | ***S. purpuratus genome**** | ***P. miniata genome**** | ***Secreted (SignalP)^2^*** | **Predicted GO terms (Predict Protein^3^)** | **Conserved domains (CDART/PFAM^4^)** |
| --- | --- | --- | --- | --- | --- | --- | --- | --- | --- |
| *Afi_tr31926* | No hit | N/A | *O. brevispinum*^♯^ | 5E-05 | no | no | yes | Binding, catalytic activity | N/A |
| *Afi_tr35695* | No hit | N/A | *O. brevispinum* | 2E-09 | no | no | yes | Cation binding, calcium ion binding | N/A |
| *Afi_tr45279* | PREDICTED: uncharacterized protein [S. purpuratus] | 2E-06 | *L. annulatus*  *O. brevispinum*  *A. mediterannea* | 10E-19  5E-10  6E-07 | no | yes | yes | Protein binding, antigen binding | Ig superfamily |
| *Afi_tr6206* | PREDICTED: titin-like [S. kowalevskii] | 2E-02 | *A. muricatum*  *O. brevispinum* | 2E-37  7E-34 | no | no | yes | Protein binding, IgG binding | N/A |
| *Afi_tr9107* | PREDICTED: exoenzymes regulatory protein AepA-like [Acropora digitifera] | 1E-41 | *A. muricatum*  *L. clathrata*  *O. spiculata* | 2E-68  3E-62  2E-22 | no | yes | no | Hydrolase activity, catalytic activity | Amidohydrolase domain |

* based on reciprocal blast

^♯^ colour scheme for echinoderm species: ophiuroid, asteroid, crinoid

^1^ NCBI non-redundant database - blast.ncbi.nlm.nih.gov/Blast.cgi

^2^ SignalP - cbs.dtu.dk/services/SignalP

^3^ Predict protein - predictprotein.org

^4^ CDART - ncbi.nlm.nih.gov/Structure/lexington/lexington.cgi; PFAM - pfam.xfam.org
